# Xrp1 integrates cellular stress through different control modes in *Drosophila*

**DOI:** 10.64898/2026.10.09.757274

**Authors:** Elisabeth Cazals, Cyrille Alexandre, Isabelle Gaugué, Tatsushi Igaki, Pierre Léopold, Jean-Paul Vincent, Carlos Estella, Laura Boulan

**Author notes:** Joint senior authors (Pierre Léopold, Jean-Paul Vincent, Carlos Estella, Laura Boulan).

## Abstract

Cells and tissues have the ability to respond to different stresses and maintain their homeostasis. The b-Zip transcription factor Xrp1 was found to play a pivotal role in this context in *Drosophila*. Xrp1 is required for the response to irradiation, in genetic conditions of ribosomal protein (Rp) deficiency leading to cell competition, and more generally in mediating the Integrated Stress Response (ISR). However, how these diverse stresses converge to and activate Xrp1 is still unclear. Our results demonstrate the existence of three separate mechanisms of Xrp1 induction. In response to Rp stress, a short isoform of Xrp1 is produced and requires the function of the ribosomal protein RpS12. In response to irradiation, a specific mRNA isoform is transcriptionally induced, which encodes the Xrp1-short protein isoform, in a p53-dependent manner. In response to the ISR, the long isoform of Xrp1 is translated by the derepression of a uORF present on the Xrp1 5’UTR, a mechanism that requires eIF2α phosphorylation, RpS12, and a transcriptional induction by ATF4. Therefore, our work provides a comprehensive understanding of the role of Xrp1 in different situations of cellular stress, illustrating the complex level of integration of the mechanisms controlling this central gene.

## Introduction

All living organisms are repeatedly subjected to a variety of environmental and intrinsic stresses such as for example exposure to toxins or ionising radiation, infections, nutrient deprivation and accumulation of misfolded proteins. In metazoans, these insults affect not only cell survival but also stem cell maintenance, differentiation timing and organ growth, and organismal homeostasis^1,2^. A plethora of mechanisms have therefore evolved to counter the effect of cell and tissue stress. *Drosophila melanogaster* has emerged as a powerful model to study the physiological response to different types of stress in the context of a whole organism. In *Drosophila*, as in human cells, the response to irradiation requires p53^3^. Irradiation also induces expression of *Xrp1*, which encodes a bZip-domain protein evolutionarily related to mammalian CHOP^4^. Core components of the integrated stress response (ISR)^5^, including PERK, GCN2, and ATF4 are conserved in *Drosophila* and this allowed demonstration of the ISR’s involvement in growth control, development, metabolic adaptation, and neurodegeneration^6,7^. Xrp1 was independently identified in screens aimed at identifying genes required for cell competition, a process whereby so-called *Minute* cells (heterozygous for a ribosomal protein-encoding gene; Rp^+/-^) are eliminated in genetic mosaics^8–10^. Xrp1 was also found to be required for the developmental delay induced by knocking down ribosomal protein-encoding genes solely in wing imaginal discs. This was found to be controlled by Xrp1-mediated expression of Dilp8, which in turn, indirectly, dampens ecdysone production in the ring gland^11^. Overall, a large body of evidence has identified Xrp1 as a central mediator of the response to diverse cellular stresses in *Drosophila*.

Molecular understanding of the mode of Xrp1 activation by different stress is still fragmentary and a comparative analysis of the response to diverse stresses is lacking. So far, Xrp1 upregulation after irradiation is thought to be mediated by p53-dependent transcriptional activation. By contrast, a completely different mechanism appears to be at work for the response to Rp-deficiency in the context of cell competition. A role for splicing was suggested by the identification of cell competition-abrogating point mutations that are located at an intro-exon junction of *Xrp1*^12^. Such role of splicing was recently confirmed by work on RpS12. Here again, a point mutant identified in a genetic screen for genes involved in cell competition (*RpS12^G97D^*) led to major insight ^13^. Phenotypic analysis showed this ribosomal protein differs from other ribosomal components. Recent results indicate that, in addition to being a component of ribosomes, RpS12 participates in the splicing of *Xrp1* mRNAs, leading to the expression of a short Xrp1 isoform that triggers apoptosis in *Rp^+/-^*(Minute) territories of genetic mosaics^14,15^. While these findings constitute a major advance towards understanding the response to Rp deficiency, a mechanistic understanding of Xrp1 induction in response to ISR has remained elusive. Phosphorylation of eIF2α, a hallmark of the ISR is activated in *Rp^+/-^* cells. However, PERK, and hence ISR, are not required for Rp^+/-^ cells to die in situation of cell competition^16^. It is likely therefore that ISR activates Xrp1 through a distinct mechanism.

In this paper we set out to compare how irradiation, ISR, and Rp deficiency activate Xrp1 expression in wing imaginal discs of *Drosophila*. We find that two Xrp1 protein isoforms are produced depending on the nature of the stress, and we decipher the underlying mechanisms. We show that irradiation triggers transcriptional activation of mRNAs encoding a short protein isoform. In contrast, the ISR uses an uORF choice mechanism to derepress translation of the long protein isoform from mRNAs that are already present in the unstressed tissue. In addition, Xrp1 induction by the ISR relies on further transcriptional activation by ATF4, which is itself regulated through an RpS12-dependent ORF choice. Our analysis of the response to Rp-deficiency is consistent with the suggestion that this stress triggers expression of a short protein isoform through a splicing-dependent mechanism. Our results show that Xrp1 is activated through at least three different modes, allowing it to integrate multiple stresses and respond accordingly.

## Results

### Xrp1 mediates cellular responses to three distinct classes of stress

As expected from the upregulation of Xrp1 in wing imaginal discs lacking one allele of a ribosome protein-encoding gene^8–10^, we found that RNAi-mediated knockdown of the gene encoding *RpL7* led to a marked increase of GFP fluorescence in wing imaginal discs from a knock in strain expressing Xrp1::GFP from the endogenous locus (Fig. 1A). A similar level of Xrp1::GFP expression was induced after activation of the ISR by Gadd34 knockdown (Fig. 1A-C). Xrp1::GFP was also upregulated in wing discs upon whole larva irradiation (Fig. 1D,E), as expected from the report that Xrp1 expression is activated in irradiated *Drosophila* embryos^4^. Therefore, three distinct stresses activate Xrp1 expression in wing imaginal discs.

**Figure 1.**
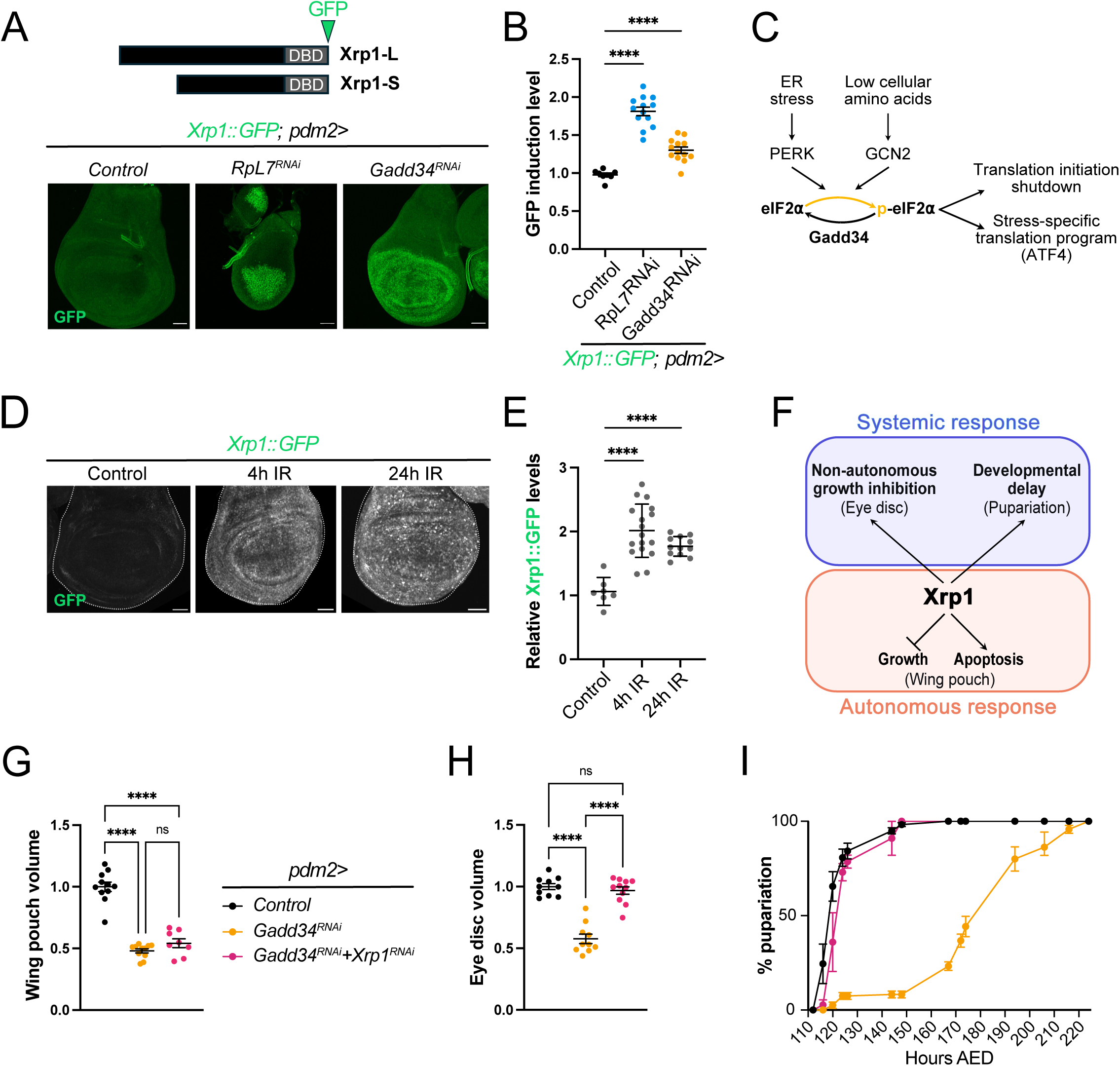
Xrp1 is activated downstream of the ISR and irradiations. A- Xrp1::GFP reporter showing the levels of both Xrp1 protein isoforms (Xrp1-L and Xrp1-S) upon expression of *RpL7-RNAi* (Rp deficiency) or *Gadd34-RNAi* (activation of the integrated stress response) under the control of the wing pouch-specific *pdm2-Gal4* driver. Maximum projections of representative wing discs are shown. Scale bar is 50μm. B- Quantification of GFP induction levels in the wing pouch (measured as the pouch/notum ratio) for the genotypes presented in A. ****p < 0.0001, ANOVA. Error bars are SEM. C- Schematic representation of the Integrated Stress Response (ISR), which is mediated by the phosphorylation of the translation initiation factor eIF2α and the transcription factor ATF4. The Gadd34 phosphatase is a key negative regulator of this pathway, counterbalancing eIF2α phosphorylation. D- Xrp1::GFP reporter showing the levels of both Xrp1 isoforms, 4 and 24 hours after irradiation (IR). Maximum projections of representative wing imaginal discs are shown. Scale bar is 50μm. E- Quantification of GFP induction levels in the wing disc, relative to the mean value of the control, for the genotypes presented in D. ****p < 0.0001, ANOVA. Error bars are SEM. F- Scheme representing Xrp1 functions as a stress response transcription factor. Upon a local stress, Xrp1 triggers an autonomous response, inducing apoptosis and growth impairment. Additionally, Xrp1 activation induces a systemic response, characterized by a non-autonomous growth inhibition, as well as a delayed developmental transition. G, H, I: Autonomous and systemic effects of ISR activation in the wing pouch (*Gadd34-RNAi*) and interaction with *Xrp1-RNAi*. The genotypes are common to the three panels: G- Tissue autonomous effect measured by wing pouch volume relative to the mean value of the control genotype *pdm2>+*. ****p < 0.0001 and ns = not significant, ANOVA. Error bars are SEM. H- Non-autonomous effect on remote tissue growth measured by eye disc volume relative to the mean value of the control genotype *pdm2>+*. ****p < 0.0001 and ns = not significant, ANOVA. Error bars are SEM. I- Systemic effect on developmental delay measured by pupariation curves displaying the percentage of animals having pupariated at the indicated time points for the indicated genotypes. Error bars are SEM. AED: After egg deposition.

Xrp1 is known to be required for apoptosis under conditions of ribosomal protein (Rp) deficiency. In addition, as we have previously shown, local Xrp1 induction triggers a systemic response (Fig. 1F) through transcriptional activation of *dilp8*. Dilp8 suppresses ecdysone synthesis, thereby delaying pupariation and inhibiting the growth of other discs ^17–19^. Similarly, activation of the ISR in the wing pouch (*pdm2>Gadd34^RNAi^*), which autonomously reduces tissue size, delays pupariation and remotely inhibits growth, e.g., in the eye disc (Fig. 1G-I). The systemic effects of ISR activation are mediated by Xrp1 since they are suppressed by co-expression of a transgene expressing RNAi against Xrp1 (*pdm2>Xrp1^RNAi^+Gadd34^RNAi^*) (Fig. 1G-I). Unlike Rp deficiency and ISR activation, gamma irradiation does not require Xrp1 to trigger apoptosis. Instead, irradiation-induced apoptosis depends on p53, as it is suppressed in a p53 mutant background^20^ (Suppl. Fig. 1A,B). Moreover, consistent with an earlier report showing that irradiation-induced developmental delay is mediated by p53 ^21^, this developmental delay also appear to be independent of Xrp1, as it still occurs in a *Xrp1* mutant background (Suppl. Fig. 1C).

Altogether, these experiments establish that Xrp1 is activated by various stresses and contributes to tissue autonomous and non-autonomous responses. We next set out to investigate how diverse stresses trigger Xrp1 expression.

### Different stresses activate distinct Xrp1 isoforms

The *Xrp1* locus encodes two protein isoforms, Xrp1-long (Xrp1-L) and Xrp1-short (Xrp1-S), which are generated by several transcription start sites and/or splicing variants (Fig. 2A). We considered the possibility that the structural complexity of the *Xrp1* locus could enable several regulatory mechanisms, thus explaining activation by diverse inputs. We first investigated *Xrp1* regulation at the transcriptional level by designing transcript-specific primer pairs to evaluate the induction levels of each transcript in different stress conditions (see Experimental Procedures and Fig. 2A). To assess the effect of Rp deficiency and ISR activation, relevant RNAi transgenes were transiently induced using with the *shine-gal4* optogenetic system ^22^ (Suppl. Fig. 2A). Knockdown of either *RpS3* or *RpL7* led to the specific induction of mRNA isoforms encoding Xrp1-S, with relatively modest induction of the Xrp1-L-encoding transcripts (A, G, B, and F) (Fig. 2B). In contrast, Gadd34 knockdown led to robust upregulation of the Xrp1-L-encoding isoform B and the Xrp1-S isoform D (Fig. 2C). Irradiation produced yet a distinct response with induction of mRNA isoform D, encoding Xrp1-S (Fig. 2D).

**Figure 2.**
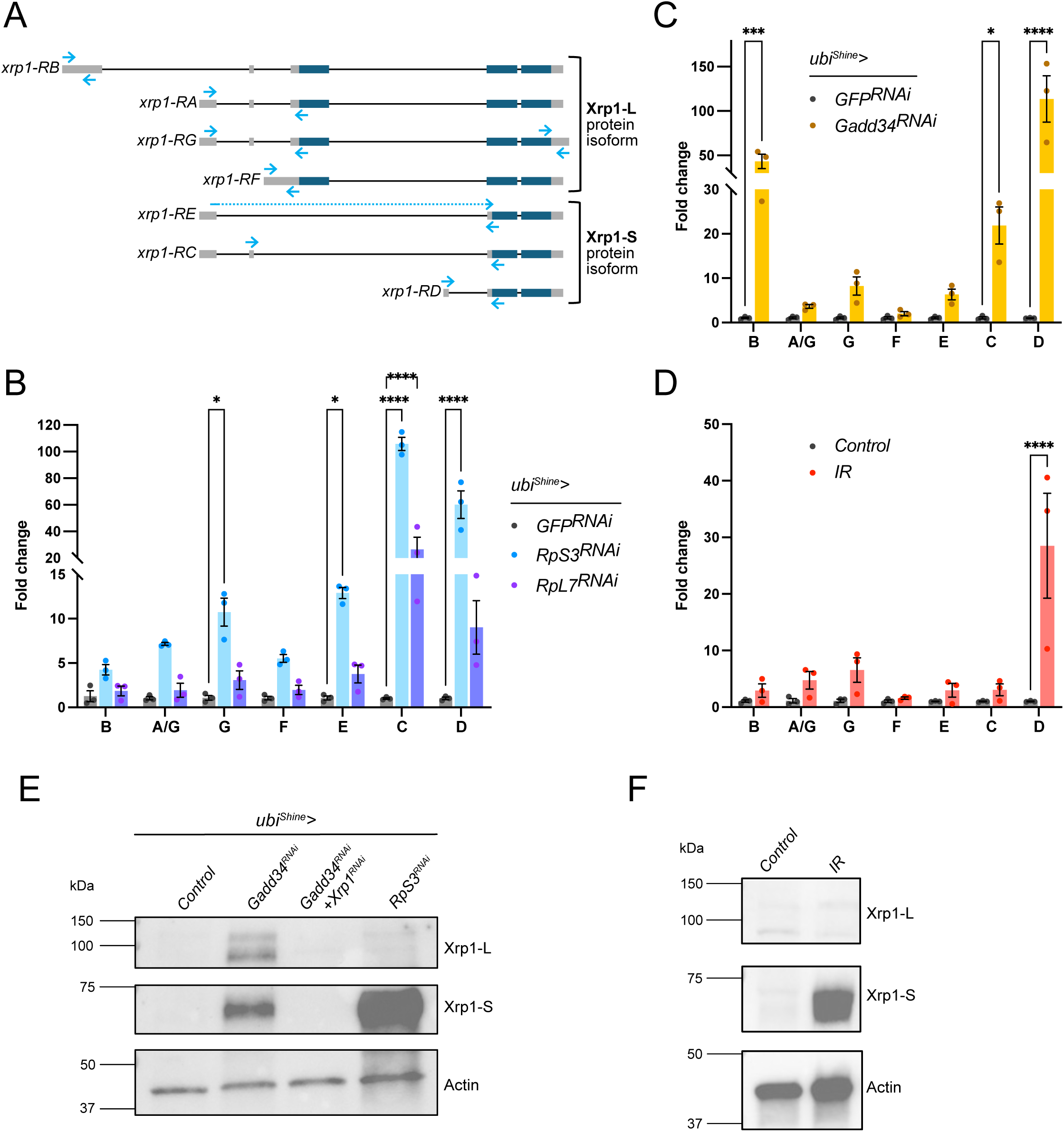
Different Xrp1 mRNA and protein isoforms are induced depending on the cellular stress context. A- Scheme representing Xrp1 transcripts. Boxes represent exons (grey are UTRs and dark blue are coding sequences). The four top mRNA isoforms give rise to Xrp1-L protein isoform, while the three bottom ones translate into Xrp1-S isoform. Light blue arrows represent the approximate localization of the primer sets used to detect each transcript by RT-qPCR (see experimental procedures). B- Relative expression levels of *Xrp1* transcripts in dissected wing discs upon Rp deficiency (*RpS3* and *RpL7* downregulation), measured by RT-qPCR and expressed as the ratio with the control genotype (*ubi^Shine^>GFP-RNAi*). ****p < 0.0001, *p = 0,1, ANOVA. Only statistically significant differences are shown. Error bars are SEM. C- Relative expression levels of *Xrp1* transcripts in dissected wing discs upon induction of the ISR, measured by RT-qPCR and expressed as the ratio with the control genotype (*ubi^Shine^>GFP-RNAi*). ****p < 0.0001, ***p= 0,001, *p<0,1, ANOVA. Only statistically significant differences are shown. Error bars are SEM. D- Relative expression levels of *Xrp1* transcripts in dissected wing discs upon irradiation, measured by RT-qPCR and expressed as the ratio with the non-irradiated control. ****p < 0.0001, ANOVA. Only statistically significant differences are shown. Error bars are SEM. E- Immunoblots of wing imaginal discs collected from L3 larvae subjected to ISR induction (*Gadd34-RNAi*) or Rp deficiency (*RpS3-RNAi*). Samples were probed with antibodies to Xrp1 and actin. Molecular weights are indicated to the left. Note that the two bands corresponding to Xrp1-S and Xrp1-L in *Gadd34-RNAi* conditions disappear upon simultaneous expression of *Xrp1-RNAi*. F- Immunoblots of wing imaginal discs from L3 larvae collected 24 hours after irradiation and probed with antibodies to Xrp1 and actin. Molecular weights are indicated to the left.

We next characterized the protein isoforms produced in each context by Western blot analysis on dissected wing discs. Upon ISR induction, two bands were observed, corresponding to the short (75kDa) and the long (100kDa) isoforms (Fig. 2E, see also^15^). By contrast, only the 75kDa band, corresponding to the short isoform, was detected upon *RpS3* knockdown (Fig. 2E), in accordance with recent analysis of *Minute* mutant discs^14,15^. Irradiation also produced only the short isoform (Fig. 2F). This observation was confirmed *in situ* by comparing two knock-in alleles: while Xrp1::GFP described above (GFP at the C-terminus, hence tagging both Xrp1-L and Xrp1-S) was activated by irradiation (Fig. 1D,E), an allele expressing V5::Xrp1-L (V5 at the N-terminus, hence tagging only Xrp1-L, also created for this study), was not (Suppl. Fig.2B). Altogether, these results show that irradiation induces the short isoform of Xrp1, possibly through transcriptional activation of the transcript isoform *Xrp1-D* (Fig. 2A,D).

### ATF4 is required for Xrp1 induction upon ISR activation but not Rp deficiency

Activation of the ISR is known to trigger phosphorylation of eIF2α and induction of the transcription factor ATF4 (Fig. 1C). Increased eIF2α phosphorylation has also been documented in Rp-deficient Minute heterozygotes^10,23,24^, suggesting that eIF2α phosphorylation could be a basic requirement for Xrp1 induction. To test this possibility, we first examined wing discs lacking one copy of the *RpS26* gene. As expected, immuno-reactivity against phospho-eIF2α (p-eIF2α) as well as Xrp1 levels (detected with an Xrp1::HA knock-in allele^9^) were markedly increased in *RpS2C*^+/-^ discs. However, suppression of eIF2α phosphorylation by Gadd34 overexpression did not impair Xrp1::HA induction (Suppl. Fig. 3A), suggesting that eIF2α phosphorylation is not required for Xrp1 induction by Rp deficiency. Accordingly, remote inhibition of eye disc growth and the pupariation delay triggered by *pdm2>RpS3-RNAi* were not rescued by Gadd34 overexpression nor ATF4 knockdown (Suppl. Fig. 3B-D), despite a decrease in p-eIF2α levels below baseline (Suppl. Fig. 3E).

We next compared the requirement of ATF4 in the upregulation of Xrp1 upon Rp deficiency and ISR activation. Using an ATF4::GFP knock-in strain^25,26^, we found that ATF4 protein is upregulated in *Gadd34-RNAi* but not *RpL7-RNAi* wing imaginal discs (Fig. 3A,B). This is mediated, at least in part, by translation control since a classical translational reporter comprising a dsRed flanked by *ATF4* 5′ UTR and regulatory uORFs ^27^ was induced by *Gadd34-RNAi* but not by *RpS3-RNAi* (Fig. 3C,D) or *RpL7-RNAi* (Suppl. Fig. 3F). We next tested whether ATF4 is required for Xrp1 activation by the ISR. Silencing *ATF4* suppressed the expression of all the *Xrp1* mRNA isoforms upregulated by *Gadd34-RNAi* (Fig. 3E) and prevented the accumulation of the Xrp1::GFP reporter (Fig. 3F). These results show that ATF4 exerts a major role in the upregulation of Xrp1 in response to ISR, but not in response to Rp deficiency.

**Figure 3.**
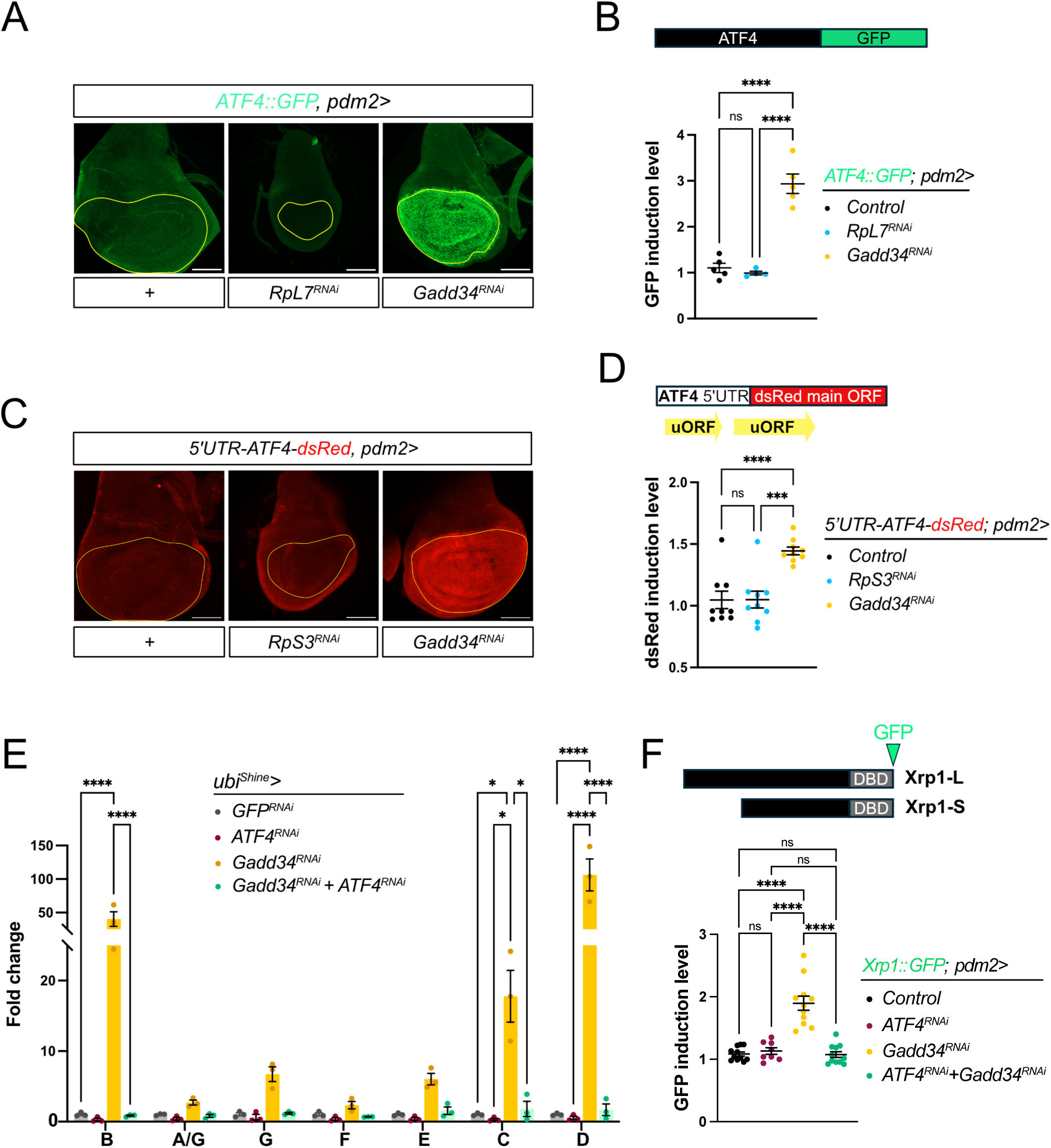
ATF4 is required for Xrp1 induction upon ISR activation, but not upon Rp deficiency. A- ATF4::GFP fusion protein reporting for ATF4 induction upon Rp deficiency (*RpL7-RNAi*) or ISR induction (*Gadd34-RNAi*) under the control of the wing pouch-specific *pdm2-Gal4* driver. The yellow line outlines the wing pouch. Maximum projections of representative wing discs are shown. Scale bar is 100μm. B- Quantification of GFP induction levels in the wing pouch (measured as the pouch/notum ratio) for the genotypes presented in A. ****p < 0.0001, and ns = not significant, ANOVA. Error bars are SEM. C- Reporter for ATF4 translation levels carrying ATF4 5’UTR containing its uORFs, and a dsRed sequence replacing ATF4 coding region (Kang et al, 2015), upon Rp deficiency (*RpS3-RNAi*) or ISR induction (*Gadd34-RNAi*). Maximum projections of representative wing discs are shown. The yellow line outlines the wing pouch. Scale bar is 100μm. D- Quantification of dsRed induction levels in the wing pouch (measured as the pouch/notum ratio) for the genotypes presented in C. ****p < 0.0001, ***p<0,001, and ns = not significant, ANOVA. Error bars are SEM. E- Relative expression levels of *Xrp1* transcripts in dissected wing discs upon induction of the ISR (*Gadd34-RNAi*) and inhibition of ATF4, measured by RT-qPCR and expressed as the ratio with the control genotype(*ubi^Shine^>GFP-RNAi*). ****p < 0.0001, *p<0,1, ANOVA. Only statistically significant differences are shown. Error bars are SEM. F- Quantification of Xrp1::GFP induction levels in the wing pouch (measured as the pouch/notum ratio) in the indicated genotypes. ****p < 0.0001, and ns = not significant, ANOVA. Error bars are SEM.

### Irradiation-induced upregulation of Xrp1-S relies on p53

We next considered the roles played by eIF2α phosphorylation in the induction of Xrp1 by irradiation. An increase in p-eIF2α immunoreactivity was detected 4 and 24 hours after irradiation (Suppl. Fig. 4A,B). However, de-phosphorylation of p-eIF2α by *Gadd34* overexpression did not prevent irradiation-induced Xrp1 accumulation or apoptosis (Suppl. Fig. 4C,D). Moreover, irradiation did not activate ATF4 (Supp. Fig. 3E), suggesting that irradiation-induced eIF2α phosphorylation does not trigger a strong cellular response.

Upon irradiation, *Xrp1-D* mRNA isoform is upregulated (Fig. 2D). Previous work detected binding of p53 in an intron close to the transcription start site of this mRNA isoform^28^, suggesting it could be a direct transcriptional target (Fig. 4A). Indeed, knocking down p53 prevented irradiation-induced upregulation of Xrp1::GFP (Fig. 4B) in accordance with the previously demonstrated role of this transcription factor in Xrp1 induction by irradiation^4^. Consistent with our Western blots showing that only Xrp1-S accumulates upon irradiation (Fig. 2F), *p53* overexpression induced Xrp1::GFP but not V5::Xrp1-L (Fig. 4C). Furthermore, we found that *p53* overexpression activated both a p-lacZ element insertion (*Xrp1-lacZ*) and a dedicated *cis* regulatory element that contains the p53 binding region of Xrp1 named *Xrp1-CRE-lacZ* (Fig. 4A,D). All together, these results indicate that p53 regulates Xrp1 most likely through transcriptional activation of the *Xrp1-D* mRNA isoform.

**Figure 4.**
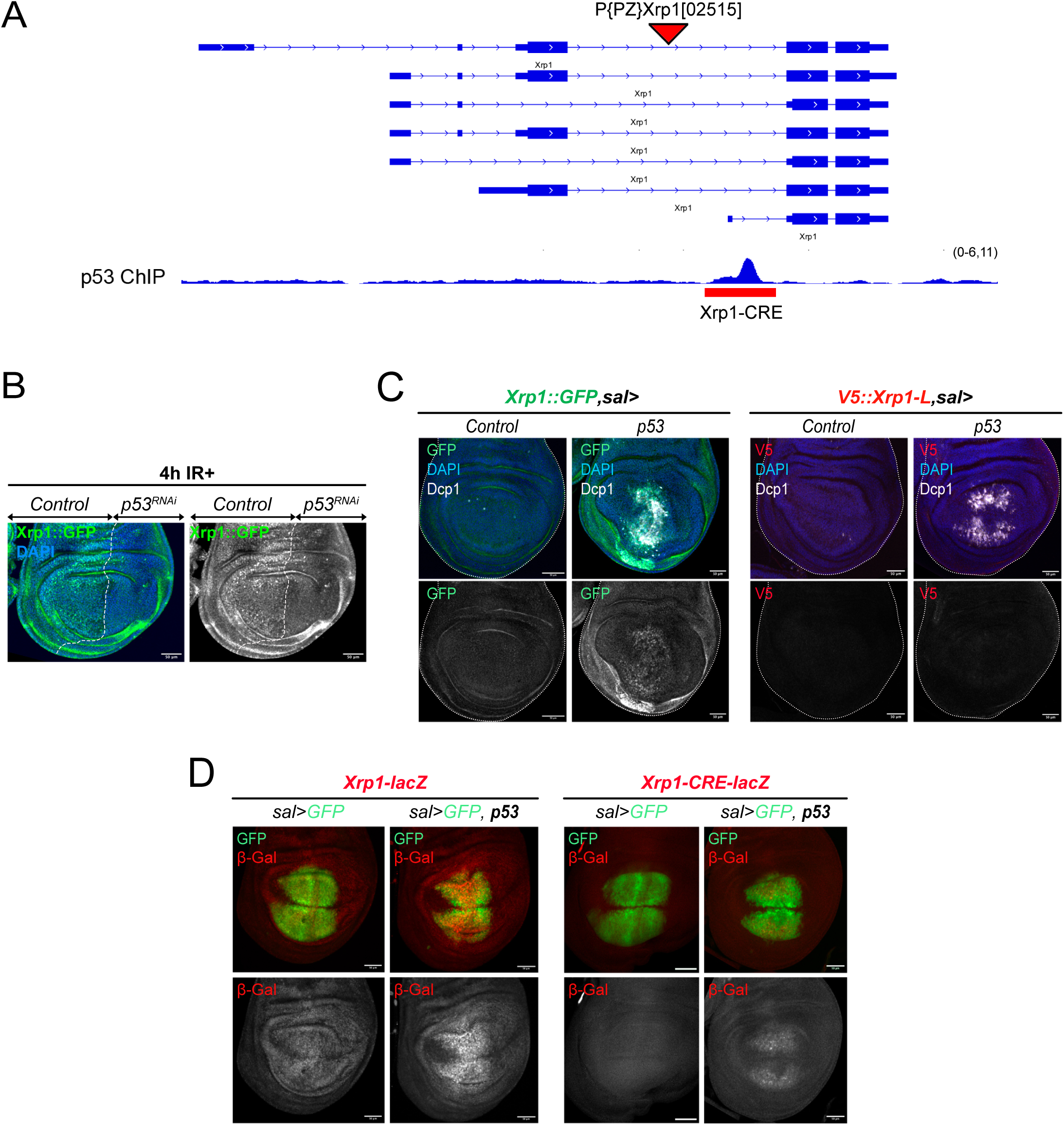
p53 is required to induce Xrp1-S upon irradiation. A. Scheme of the *Xrp1* locus together with the ChIP track for p53 obtained from (Kurtz et al, 2019), showing the p53 binding region close to the Xrp1-RD first exon. The region corresponding to the cis regulatory element (CRE) and cloned to generate the Xrp1-CRE-lacZ reporter spans the p53 binding region and its marked by a red bar. Also shown is the insertion site of a p-lacZ element in the Xrp1-lacZ reporter line (red triangle). B. Induction of the Xrp1::GFP reporter in representative wing discs expressing an RNAi targeting p53 in the posterior compartment and dissected 4 hours after irradiation. RNAi was induced for 48 hours under the control of a *hh-Gal4* driver combined with a *Gal80ts* construct for temporal control using temperature shift. Scale bar is 50μm. C. Induction of the Xrp1::GFP reporter and the V5::Xrp1-L in representative discs overexpressing p53 in the central region of the wing pouch under the control of *sal-Gal4*. Scale bar is 50μm. D. Overexpression of p53 with the *sal-Gal4* driver induces the Xrp1-lacZ transcriptional reporter, as well as the cis regulatory element (CRE) of Xrp1, where p53 binds. Scale bar is 50μm.

### Translation of the long isoform of Xrp1 is repressed by an upstream ORF

It is worth noting that Xrp1-L protein is not expressed in unstressed tissue despite abundant mRNAs coding for this isoform, especially *Xrp1-A* and *Xrp1-G* (Suppl. Fig.5A) (See also^14,15^). This apparent disconnect between RNA and protein expression is reminiscent of ATF4 regulation, which relies on upstream Open Reading Frames (uORFs)^29^. For ATF4 and other ISR effectors, translation is initiated by default at uORFs, preventing translation of the coding ORF under non-stressed conditions. Upon ISR activation, these uORFs are bypassed by the ribosome, allowing translation of the coding ORF through an “ORF-choice” mechanism^30^. A 5’UTR structure similar to that of ATF4 has been noted in the 5’ UTR of the *Xrp1* transcripts that encode Xrp1-L (Supp. Fig. 5B)^31^. In particular, all mRNA isoforms coding for Xrp1-L share a uORF overlapping with Xrp1-L first exon, suggesting that this uORF could have inhibitory function on the translation of Xrp1-L , as suggested by *in vitro* studies^15, 32^To test this *in vivo*, we deleted a 95 bp region encompassing the ATG of the uORF using a tissue-specific CRISPR-Cas9 approach (Fig. 5A). *Xrp1::GFP* flies ubiquitously expressing the gRNA pair targeting the uORF were crossed with flies expressing Cas9 in the wing pouch (*nub-GAL4>UAS-Cas9*). Robust GFP expression was detected in wing discs expressing both the Cas9 and the gRNAs but not in the not in the absence of (Fig. 5B,C). Therefore, it appears that inactivation of Xrp1’s uORF derepresses translation of the main ORF, leading to Xrp1::GFP protein accumulation. Note that significant apoptosis appeared in this background (Fig. 5B,D), suggesting that the pool of Xrp1 produced in this context is functional. The same editing strategy was conducted in a strain expressing, from the endogenous locus, N-terminally tagged Xrp1 (V5::Xrp1-L) to specifically mark the long isoform. V5 immunoreactivity was detected in the wing pouch upon introduction of the gRNA-expressing and *nub>Cas9* transgenes (Fig. 5E,F), confirming that Xrp1-L is upregulated and underscoring the importance of this regulatory uORF *in vivo* (see Fig. 2A). Since Cas9 activation leads to non-specific DNA damage, which in turn could induce Xrp1, we tested whether control gRNAs targeting the *white* gene (gRNA_white_) had any impact on Xrp1 expression. Flies ubiquitously expressing Cas9 and gRNA_white_ exhibited a mosaic of red and white ommatidia, confirming targeting of *white* (Supp. Fig. 5C). Targeting *white* in the wing pouch did increase the level of apoptosis but not to the same extent as targeting *Xrp1’s* uORFs (Supp. Fig. 5D) and Xrp1::GFP was not induced by *white* targeting (Supp. Fig. 5E). Moreover, targeting *Xrp1* uORF, but not *white,* led to increased DNA damage, as indicated by immunoreactivity against γH2Av, (Supp. Fig. 5F). This was suppressed by concomitant expression of Xrp1-RNAi, suggesting that DNA damage is a consequence of Xrp1’s pro-apoptotic activity (Supp. Fig. 5G). As an additional test of the inhibitory role of the common uORF on Xrp1-L translation, we generated a reporter based on a 15kb genomic fragment previously shown to rescue Xrp1 mutations^8^. In this fragment, the uORF were deleted, and the region encoding the DNA-binding domain was replaced by GFP-encoding sequences to both tag the product and inactivate Xrp1 (thus preventing lethality from ectopic expression). The resulting transgenic larvae, but not control transgenics with the uORF left in place, expressed GFP (Fig. 5G). Overall, the above data indicate that Xrp1’s uORF normally prevents expression of Xrp1-L in unstressed imaginal discs despite the presence of abundant mRNA. They also suggest that, as for ATF4, the ISR could derepress this mechanism to allow translation of Xrp1-L. Thus, Xrp1-L would be controlled at two levels: transcriptionally by ATF4, and translationally through an “ORF-choice” mechanism.

**Figure 5.**
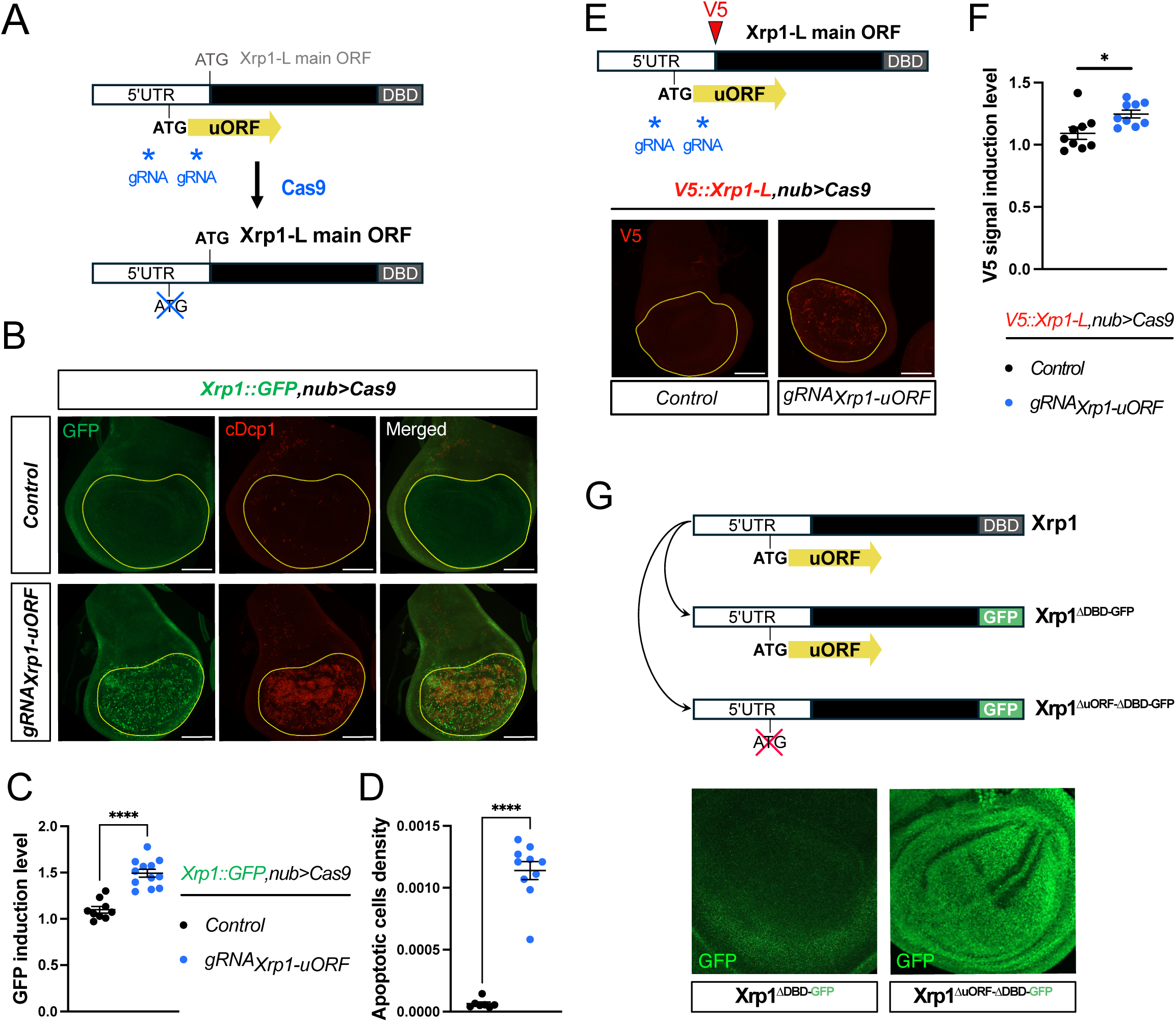
An upstream ORF represses Xrp1-L. A- Scheme of the strategy to mutate the common uORF of Xrp1 in a tissue-specific manner. A pair of guide RNAs (gRNA) that recognizes the flanking region of the uORF ATG was designed to induce a deletion of this small region upon Cas9 expression. The blue stars mark the approximate locations of the gRNAs. B- Induction of Xrp1::GFP and apoptosis upon deletion of Xrp1 uORF in otherwise non-stressed tissues using wing pouch-specific expression of the Cas9 with the *nub-Gal4* driver. Maximum projections of representative wing discs are shown, stained for GFP and cDcp1, and yellow lines outline the wing pouch region. Scale bar is 100 μm. C- Quantification of GFP induction levels in the wing pouch (measured as the pouch/notum ratio) for the genotypes presented in B. ****p < 0.0001, t-test. Error bars are SEM. D- Quantification of apoptotic cell density in the wing pouch for the genotypes presented in B. ****p < 0.0001, t-test. Error bars are SEM. E- Induction of the V5::Xrp1-L reporter upon deletion of Xrp1 uORF in otherwise non-stressed tissues using wing pouch-specific expression of the Cas9 with the *nub-Gal4* driver. Maximum projections of representative wing discs are shown. Yellow lines outline the wing pouch region. Scale bar is 100 μm. F- Quantification of V5 signal induction levels in the wing pouch (measured as the pouch/notum ratio) for the genotypes presented in E. *p < 0.1, t-test. Error bars are SEM. G- Top: Strategy to mutate the ATG of the common uORF while replacing Xrp1 DNA binding domain with GFP to prevent lethality. Bottom: Maximum projections of representative wing discs showing the expression levels of the reporters. Note that removing the uORF is sufficient to trigger the expression of the reporter.

### RpS12 is involved in the induction of Xrp1 by ISR

Previous work showed that the ribosomal protein RpS12 is required for the activation of Xrp1 expression in response to ribosomal deficiency^13,19,33^. We used the Xrp1::GFP reporter to test whether RpS12 also contributes to Xrp1 expression in response to ISR or irradiation. First, we confirmed that RpS12 knockdown suppressed Xrp1::GFP upregulation induced by RpL7 knockdown (Rp deficiency), even though, on its own, it led to only a mild increase (Fig. 6A,B). Similarly, RpS12-RNAi suppressed Gadd34-RNAi-induced Xrp1::GFP accumulation (Fig. 6A,B). In addition, it alleviated apoptosis (Suppl. Fig. 6A), as well as non-autonomous growth reduction and developmental delay associated with Gadd34 knockdown (ISR) (Suppl. Fig. 6B-D). We conclude that RpS12 is required for tissue-autonomous Xrp1 induction and the systemic responses to the ISR. Finally, we tested a potential role of RpS12 in irradiation-induced Xrp1-S expression. A point mutation (*RpS12^G97D^*) that prevents Rp deficiency-induced Xrp1 upregulation without interfering with ribosomal function was previously identified in a genetic screen^33^. Recently, this mutation was shown to affect the splicing that leads to the production of Xrp1-S in Minute heterozygotes^14,15^. We observed that irradiation induces strong Xrp1-S protein accumulation even in an *RpS12^G97D^* mutant background (Fig. 6C), similar to irradiation in a control genotype (Fig. 2F). This suggests that the transcriptional induction of Xrp1-S by irradiation does not relies on the splicing function of RpS12 and further supports the notion that distinct mechanisms activate Xrp1 depending on the type of stress.

**Figure 6.**
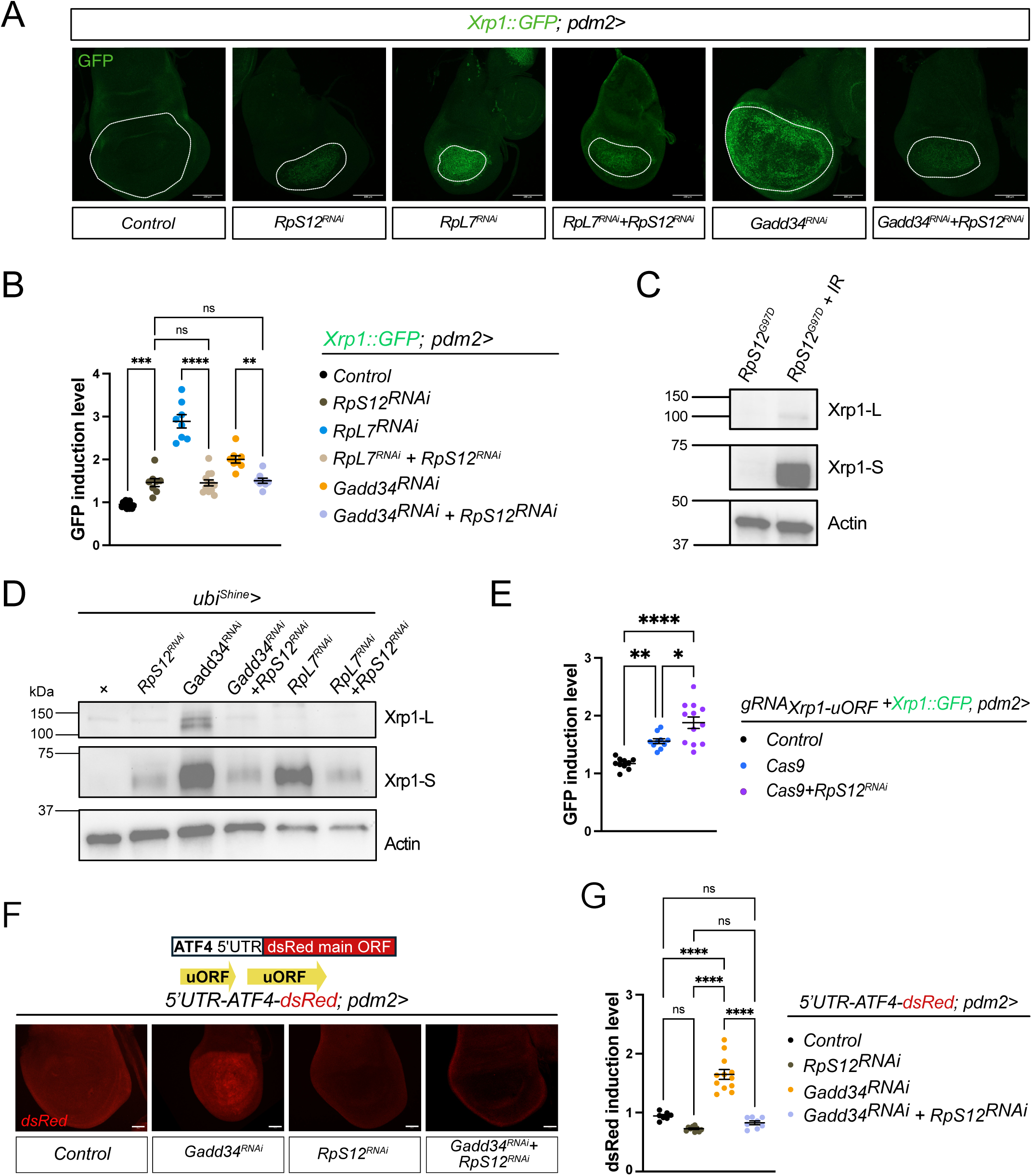
RpS12 is required for the translation of transcripts regulated by uORFs. A- Representative wing discs showing the levels of the Xrp1::GFP reporter upon wing-pouch specific inhibition of RpS12 (*RpS12-RNAi*) in the context of ISR activation (*Gadd34-RNAi*) or Rp deficiency (*RpL7-RNAi*). Maximum projections are shown, and the white line outlines the wing pouch. Scale bar is 100μm. B- Quantification of GFP induction levels in the wing pouch (measured as the pouch/notum ratio) for the genotypes presented in A. ****p < 0.0001, ***p < 0.001, **p < 0.01, and ns = not significant, ANOVA. Error bars are SEM. C- Immunoblots of wing imaginal discs from larvae with an *RpS12^G97D^* mutant background. Irradiated samples were collected 24 hours after irradiation and probed with antibodies to Xrp1 and actin. Molecular weights are indicated to the left. D- Immunoblots of wing imaginal discs from larvae upon RpS12 inhibition (*RpS12-RNAi*) in the context of ISR induction (*Gadd34-RNAi*) or Rp deficiency (*RpL7-RNAi*). Samples were probed with antibodies to Xrp1 and actin. Molecular weights are indicated to the left. E- Quantification of Xrp1::GFP induction levels in the wing pouch (measured as the pouch/notum ratio) upon derepression caused by the mutation of Xrp1 uORF and the concomitant downregulation of RpS12. ***p < 0.001, **p < 0.01, and *p < 0.1, ANOVA. Error bars are SEM. F- Reporter for ATF4 translation levels upon ISR induction (*Gadd34-RNAi*) and RpS12 knockdown in the wing pouch (*pdm2-Gal4*). Maximum projections of representative wing discs stained for dsRed are shown. Scale bar is 50μm. G- Quantification of dsRed induction levels in the wing pouch (measured as the pouch/notum ratio) for the genotypes presented in F. ****p < 0.0001, and ns = not significant, ANOVA. Error bars are SEM.

### RpS12 is required for uORF-regulated translation

While Rp deficiency only activates Xrp1-S, both the long and short isoforms are expressed upon ISR (see Fig. 2E) and Western blot analysis showed that RpS12 knockdown prevents the production of both isoforms (Fig. 6D). The underlying mechanism is unlikely to involve splicing since Xrp1-L-encoding transcripts are not subjected to splicing. Of potential relevance, we found that activation of Xrp1::GFP by deletion of the uORF is unaffected by RpS12 knockdown (Fig. 6E), suggesting that reduction of ISR-induced Xrp1 expression by RpS12-RNAi is not due to a general decrease in translation efficiency. Therefore, it appears that, in addition to its role in controlling *Xrp1* mRNA splicing, RpS12 could be needed for uORF-mediated control on Xrp1 translation. To test this hypothesis, we turned to ATF4, the expression of which is controlled by uORFs. ISR-induced activation of an ATF4 translation reporter comprising the 5’UTR of *ATF4* upstream of DNA encoding dsRed^27^ was found to be inhibited by RpS12 knockdown, confirming that RpS12 contributes to ORF selection (Fig. 6F,G). Moreover, knocking down RpS12 did not prevent remote growth inhibition or pupariation delay upon ectopic ATF4 production from a transgene lacking the 5’ UTR (Supp. Fig. 6B-D). Therefore, removal of uORF in both *ATF4* and *Xrp1-L* transcripts renders their translation insensitive to RpS12. Altogether, these observations suggest that RpS12 contributes to the translational control of mRNAs containing uORFs.

## Discussion

Xrp1 has been shown to be activated by diverse stresses such as Rp deficiency, the integrated stress response and irradiation. The underlying mechanisms have so far been studied in isolation. Here we took a comparative approach to identify stress-specific regulatory features and shared mechanisms, with the aim of understanding how the response to stresses is coordinated. The three classes of stresses trigger expression of Xrp1, which in turn mediates autonomous (e.g. apoptosis) and non-autonomous (e.g. developmental delay) responses. We found that, depending on the nature of the stress, multiple mRNAs and two protein isoforms, Xrp1-Short and Xrp1-Long, are produced. For each stress, we found that various cellular processes contribute differentially to Xrp1 upregulation (Table 1). This allows Xrp1 to integrate sensing of a wide range stresses, presumably optimizing the overall response and organismal homeostasis.

**Table 1.** Summary of the different modes of Xrp1 regulation depending on the cellular stress.

|  | RP DEFICIENCY<br>(Minute or Rp-RNAi) | IRRADIATION | ISR (Gadd34-RNAi) |
| --- | --- | --- | --- |
| Protein Isoforms | Short only | Short only | Short and long |
| P53-dependent? | NO | YES | Not tested |
| Sensitive<br>to *RpS12*<sup>G97D</sup>? | YES | NO | Not tested |
| Alternative splicing<br>dependent? | YES | NO | Presumed |
| Regulated by ORF<br>choice? | NO | NO | YES (Xrp1-Long) |
| Dependent on eIF2α<br>phosphorylation? | NO | NO | YES |
| ATF4-dependent? | NO | Unlikely since<br>ATF4-GFP not<br>induced | YES. All mRNAs are<br>upregulated by ATF4 |

We first compare Rp deficiency and irradiation, which both induce expression of Xrp1-S protein only. Rp deficiency leads to Xrp1-S expression through RpS12-dependent *Xrp1* mRNA splicing, a process that is affected by the *RpS12^G97D^* mutation^14,15^, but not p53^34^. In contrast, the production of Xrp1-S by irradiation is unaffected by the *RpS12^G97D^* mutation but depends on p53-dependent transcription of *Xrp1-D*, which encodes Xrp1-S. Therefore, even though irradiation and Rp deficiency both trigger production of Xrp1-S, they differ in their requirement for alternative splicing and p53-mediated transcriptional activation.

Since the ISR triggers production of both Xrp1-S and Xrp1-L, it must rely on yet another mechanism. The observation that Xrp1-L is not expressed in unstressed condition despite the presence of encoding RNAs points to a post-transcriptional regulation. We noted that *Xrp1*’s 5’UTR resembles that of *ATF4* in that it harbours uORFs. In normal tissues, *ATF4*’s uORFs are thought to dampen translation by preventing ribosomes from reaching the initiating codon of the main ORF. Upon activation of the ISR, eIF2α is phosphorylated, allowing ribosome to reach the downstream initiation site and translate the full length ATF4 protein. Since Gadd34-RNAi, which boosts eIF2α phosphorylation, triggers Xrp1-L expression, we considered the possibility that uORFs could similarly control Xrp1 long translation. Accordingly, deletion of *Xrp1*’s uORFs, either at the locus or in a transgenic reporter, led to increased translation from the coding ATG, indicating that the ISR directly controls Xrp1-L translation. We suggest therefore that the presence of uORFs normally prevent translation from mRNAs that encode the long isoform. The response to Rp deficiency or irradiation does not require eIF2α, as expected since only the short form is produced. Surprisingly however, phosphorylation of eIF2α upon Rp deficiency or irradiation, does not suffice to trigger Xrp1-L translation, perhaps because it is lower than the level of phosphorylation induced by ISR and not sufficient to alter uORF choice. In addition to activating Xrp1-L, ISR also triggers the production of Xrp1-S, which is not directly activated by a uORF choice mechanism. We provide evidence that this is mediated by ATF4-dependent transcription of Xrp1-S -encoding mRNA. Thus, the ISR (p-eIF2α) boosts Xrp1 expression through two routes, directly by translational de-repression of Xrp1-L and indirectly by enabling translation of the transcriptional activator ATF4.

RpS12 holds a special place among ribosomal proteins. As mentioned above, a point mutation in *RpS12* (*RpS12^G97D^*) suppresses apoptosis of cells deficient for any other *Rp* gene. Recent work has shown that this activity is mediated by an extra-ribosomal function whereby RpS12 would moonlight as a splicing factor that excises the uORFs-containing exon of Xrp1, thus causing translation to produce only Xrp1-S^14,15^. We confirmed by RNAi that RpS12 is required for the production of Xrp1-S in response to Rp deficiency. Unexpectedly however, RpS12 was also found to be required for the translation of Xrp1-L, and ATF4. This is surprising since expression of neither protein requires alternative splicing. For both Xrp1 and ATF4, epistasis analysis places RpS12 within- or upstream of-ORF choice. We propose therefore that full RpS12 activity could be required for proper regulation of uORF choice. Structural analysis reveals that RpS12 is located at the tip of the 40S beak, a prominent structure of the small subunit that plays major roles in translation, particularly its initiation^35–37^. We propose that a “scanning malfunction” of the ribosome could compromise ORF choice in *RpS12* knockdown conditions. These results echo a recent study proposing that a regulatory network of ribosomal proteins and components of translation initiation complexes controls ORF-choice dependent ATF4 translation^38^. Future work may help clarify the role of RpS12 and possibly other components of the ribosomal subunits in the ORF choice mechanism.

## RESOURCE AVAILABILITY

### Lead contact

Requests for further information and resources should be directed to and will be fulfilled by the lead contact, Laura Boulan.

### Materials availability

All reagents generated in this study are available from the lead contact.

### Data and code availability

This study did not generate data sets or codes.

## ACKNOWLEDGEMENTS

We thank all lab members for insightful discussions, Carine Ganem-Elbaz for technical help and Paula Santabárbara-Ruiz for supervision. We acknowledge the Bloomington Stock Center and the Vienna Drosophila RNAi Center for fly stocks; the PICT-IBiSA@BDD light microscopy facility of Institut Curie (member of France-Bioimaging ANR-10INBS-04); the imaging facility MRI (member of France-BioImaging supported by ANR-24-INBS-0005 FBI BIOGEN) and the Drosophila Facility from BioCampus Montpellier. We thank T. Igaki for the generous gift of anti-Xrp1 monoclonal antibodies, as described in^15^. This work was funded by La Ligue contre le Cancer (PhD fellowship to EC), an European Research Council Advanced grant no. 694677 (PL), two ANR-11-LABX-0044, ANR-10-IDEX-000102 (PL), a Human Frontier Science Program RGP0031/2020 (PL). J-PV and CA were funded by an MRC project grant (MR/W017881/1) and core funding from the Francis Crick Institute (FC001204). CE was funded by Ministerio de Ciencia e Innovación PID2024-160216NB-I00. LB was funded by CRBM, the ATIP-Avenir Program from CNRS and the I-SITE Excellence Program of the University of Montpellier, under the Investissements France 2030.

## AUTHOR CONTRIBUTIONS

Conceptualization, E.C., L.B., J-P.V., C.E., C.A., I.G. and P.L.; Methodology, E.C., L.B., J-P.V., C.E., C.A., I.G., T.I. and P.L.; Investigation, E.C., L.B., C.E., I.G. and C.A.; Writing – Original Draft, E.C.; Writing – Review & Editing, E.C., J-P.V., C.E., P.L. and L.B.; Funding Acquisition, P.L. J-P.V., C.E. and L.B.; Supervision, J-P.V., P.L., C.E. and L.B.

## DECLARATION OF GENERATIVE AI AND AI-ASSISTED TECHNOLOGIES IN THE WRITING PROCESS

During the preparation of this work, the authors used AI-assisted technologies in order to improve the readability and language of the manuscript. After using this tool/service, the authors reviewed and edited the content as needed and take full responsibility for the content of the published article.

## SUPPLEMENTAL INFORMATION

Supplementary figures 1-7 and Table1

## EXPERIMENTAL PROCEDURES

### Drosophila melanogaster husbandry

The flies were raised at 25°C (unless otherwise indicated in the figure legends or experimental procedures) on a medium composed of 7.5g agar, 35g of organic wheat flour, 50g of yeast, 55g of sugar, 25mL of methyl and 4mL of propionic acid per liter of water.

### Drosophila lines

*p53*^5a14^ (BDSC#6815), *UAS-6xMyc:p53-A*^39^, *UAS-p53-RNAi* (BDSC#29351), *UAS-RpL7-RNAi* (BDSC#34600), *UAS-RpS12-RNAi* (VDRC#109381), *UAS-RpS3-RNAi* (VDRC#37741), *UAS-ATF4-RNAi* (VDRC#109014), *RpS2C^KO^* ^23^, *UAS-dATF4/TM6B* (BDSC#81655), *UAS-Gadd34-RNAi* (VDRC#107545), *UAS-Gadd34-RNAi* (VDRC#15238), *UAS-Gadd34* (BDSC#76250), *UAS-Xrp1-RNAi* (BDSC#34521), *V5::Xrp1-L* (This study), *Xrp1::GFP/TM6B* (This study), *Xrp1-lacZ* (BDSC#11569), *Xrp1-CRE-lacZ* (This study), *tub>5’UTR-dATF4-dsRed/Cyo*^27^, *dATF4::gfp* (BDSC#59608), *nub-Gal4/Cyo;UAS-CasS/TMCB* (BDSC#67086), *U6:3>2xgRNA-Xrp1-ATG-uORF/Cyo* (*qRNA_Xrp1-uORF_*; This study), *ubi>gRNA_white_* (BDSC#79543), *Xrp1^ΔDBD-GFP^/TMCB* (This study), *Xrp1^ΔuORF^*^2^*^−ΔDBD-GFP^/TM3,sb* (This study), *w*^1118^ (BDSC#3605), *pdm2^R11F^*^02^*-Gal4* (BDSC#49828), *hh-Gal4* (Flybase #FBti0017278), *sal-Gal4* (FlyBase FBtp0021755), *ubi ^Shine^-Gal4* ^40^.

### Generation of Xrp1::GFP fly line

We generated a “donor vector” (backbone kindly provided by Y. Bellaiche), which carries the sequence coding for the GFP flanked by *Xrp1* homology regions (HR) to allow its insertion in 3’ of *Xrp1.* The GFP thus tags both the short and long isoforms of Xrp1 in C-Ter. The “carrier vector” (pCFD5, addgene #73914) contains sequences coding for gRNAs, which will mediate the recognition of the *Xrp1* locus by the Cas9, triggering the insertion of the tag through homologous recombination with the “donor vector”. The two plasmids were injected together into a fly strain expressing Cas9 by the BestGene company. Transformant offspring were then selected, balanced, and tested for Xrp1::GFP expression.

<u>gRNA1:</u> TAGCCATAGAATAGAAATTG <u>gRNA2:</u> ACGACGATAGGTACTATGAC

<u>HR1:</u> GCCTTGTGGAGGACGTCAGCG <u>HR2:</u> TACGTTAAGCAGGAGCAGGAC

### Cloning of Xrp1-CRE in a lacZ reporter vector

The sequence containing the p53 binding peak at the Xrp1 locus was PCR amplified and cloned between AvrII and Stu1 sites in the attB-hs43-nuc-lacZ plasmid vector. The Xrp1-CRE-lacZ construct was inserted into the 86Fb attP site using PhiC31 integrase.

The primers used are :

Forward : CAGTCCTAGGGCATTGCGAATGCGCACTG

Reverse : CAGTAGGCCTGGCGATCCATACATAGAACACCC

### Generation of Xrp1^ΔDBD-GFP^ and Xrp1^ΔuORF-ΔDBD-GFP^

In order to avoid any deleterious effects coming from these reporters (excess Xrp1 protein), we modified a 16kb Xrp1 rescuing construct by first deleting the DNA binding domain and replacing it with eGFP. This first modified BAC (Xrp1ΔDBD-GFP) was subsequently modified in the 5’utr of Xrp1 long isoforms (A, B, F and G) by deleting the uORF to generate Xrp1ΔuORF-ΔDBD-GFP. The 2 new reporters of Xrp1 were then inserted in the attP2 site on the 3d chromosome. All details and sequences are available upon request.

### Generation of V5::Xrp1-L

For directly tagging Xrp1 with the V5 epitope, we used the protocol describe in^41^. The starting plasmid IOGFP@NTV-Pax-Cherry was modified to express the V5 epitope (IOV5@NTV-Pax-Cherry). Using CRISPR, we integrated the plasmid with a target site (agatataatattg<u>ATG</u>atccAGG) specific to the ATG of the long isoforms. This first integration generates Xrp1 mutants for the long isoforms. Subsequent removal of the Pax-CHERRY marker with hs-CRE generates an N-ter V5 tagged Xrp1 for the long isoforms A, B, F and G. All details and sequences are available upon request.

### Generation of an RpS12^GG7D^ allele marked with *white^+^*

The RpS12^G97D^ allele was generated in two steps by CRISPR-mediated homologous recombination according to^41^. A knockout allele was first generated by replacing the three coding exons, along with the 5’ and 3’ UTRs, with an *attP* landing site and a LoxP-flanked *Pax-Cherry* cassette. The resulting strain was homozygous lethal, as expected from the essential role of RpS12. The Pax-Cherry cassette was excised by Cre-mediated recombination before Phi31-mediated insertion of a *white+* RIV *attB* vector^42^ comprising all the deleted sequences (including the 5’ and 3’ UTRs) except for a mutation to change codon 97 from G to D. The resulting strain is homozygous viable confirming that the G97D mutation does not significantly interfere with ribosomal function^33^. The mutation was confirmed by sequencing of relevant PCR products obtained from genomic DNA of homozygous flies. Details and primer sequences are available upon request.

### Cas9-mediated uORF deletion

The sequence of *Xrp1-RB* was used to generate gRNAs targeting the flanking region of Xrp1 uORF. The two target sequences are below, with the PAM sequence in bold. The Cas9 induces a double-strand break 3-4 nucleotides upstream. The complementary gRNAs were cloned into the pCFD5 plasmid (Addgene #73914), which allows for the ubiquitous expression of two gRNAs. The plasmid was sent to the BestGene company and injected into embryos. Transformant offspring were then selected, balanced, and tested for proper insertion. The flies ubiquitously expressing the gRNAs were then crossed to flies expressing *UAS-Cas9* under the control of *nubbin-Gal4*.

gRNA 1 (+strand)

Selected target:

AAGCCCAAACCAGCCAAGCCAGG

gRNA 2 (-strand)

Selected target:

GATCGGTGCAGTTTACGGTTGG

### Stainings

Tissues dissected from larvae in 1x PBS at the indicated stages were fixed in 4% paraformaldehyde (ThermoScientific, #28908) diluted in PBS for 20 min at room temperature (RT), then washed in PBS containing 0.3 % Triton-X-100 (PBT), blocked in PBT containing 2% BSA, and incubated overnight with primary antibodies at 4 °C. The next day, tissues were washed, blocked, incubated with secondary antibodies (1/400) for 2 hours at RT and finally kept in Vectashield medium (Vector Labs, #H-1000-10).

The following primary antibodies were used: mouse anti-nub 1/50 (DSHB, #2D4), mouse anti-wg 1/200 (DSHB, #4D4), rat anti-Ci (1:50, DSHB #2A1), chicken anti-GFP 1/10 000 (Abcam, #13970), rabbit anti-cDcp1 1/100 (Cell Signaling, #9578), rabbit anti-V5 1/500 (Cell Signaling #13202), rabbit anti-p-eIF2*a* 1/100 (Cell Signaling, #3398), rabbit anti-dsRed 1/500 (Takara, #632496), and rabbit anti-*γ*H2Av 1/1000 (Rockland, #600-401-914).

Secondary antibodies used: Alexa Fluor™ 488 (Invitrogen, # A-11039 and # A-11008), Alexa Fluor™ 546 (Invitrogen, # A-11030, # A-11035) and Alexa Fluor™ 488 (# A-21236). Dapi was also used at 1/1000 (FISHER SCIENTIFIC S.A.S., # 62248).

Fluorescent signal quantification:

On the Fiji software, z-stack images of discs were used to perform a Maximum-projection. The pouch region was delimited using the Nubbin or Wingless signal, defining a region of interest (ROI). The fluorescent signal (mean grey value) was then measured in the preferred channel, within the ROI. Then the signal was measured outside of the ROI (in the hinge and the notum). The values for fluorescent signal were plotted as the signal in the pouch normalized by the signal in the notum for each disc.

### Larval synchronization

Crosses were incubated at 25 °C, and eggs were collected on plates made of 2% agar and 2% sucrose in PBS. The next day, larvae that had already hatched were discarded. Synchronized L1-stage animals were collected and transferred 4 hours later from the agar plates to vials containing fly food.

### Measurements of tissue volume

Wing and eye discs were mounted in glass-bottom cell culture plates (“Cellview”; Greiner Bio-One, #627861) to preserve the 3D structure. The Vectashield mounting medium was used (Vector Labs, #H-1000-10).

Confocal Z-stack images were acquired with a Zeiss LSM900 microscope. In the Imaris software, pouch volume was measured by generating a surface based on the nubbin signal. This surface was then used as a “mask” to generate a new channel corresponding to the pouch in the DAPI channel. A surface was finally constructed using this channel, and the pouch volume was measured. The volume of the eye disc was measured by using the surface tool based on the DAPI signal. Finally, the data were normalized with the mean volume of the controls.

### Gamma Irradiation

Third instar larvae were irradiated at 40 Gys in a Gamma rays IBL-437C Cs137 irradiator. Samples were collected 4 or 24 hours after irradiation.

### Optogenetic induction of UAS transgenes expression

To prevent lethality or organ size reduction that would impair experimental procedures, we took advantage of the optogenetic system *Shine-Gal4*^40^, which allows for tight regulation of UAS construct expression. Larvae were reared at 29°C in the dark for 72 hours after egg deposition. They were then incubated for 24 hours in the light and at 29°C, inducing RNAi expression. Samples were then collected at 96 hours post egg-laying.

### Apoptosis and DNA damage quantification

For most experiments, the Imaris software was used to segment pouches based on the nubbin signal, allowing for pouch volume measurement as described above. cDcp1 and *γ*H2Av-positive cells were counted within the pouch segments by using the spot function of the Imaris software. The density of each signal was obtained by dividing the number of spots by the volume of the pouch. Apoptosis quantification in Fig.1I: Z-maximal intensity projection was generated for each image, and a high-intensity threshold was adjusted for each image. The percentage of staining covered in the region of interest was then calculated.

### Pupariation curve

Three to four groups of 20-30 synchronized larvae of each genotype are placed in tubes and raised at 25°C. The number of new pupae is scored twice a day until all larvae pupariate. The percentage of larvae that had pupariated is then plotted as a function of time.

### Western blot

50 to 75 wing imaginal discs were dissected in ice-cold PBS containing Halt™ Protease & Phosphatase Inhibitor Cocktail and immediately frozen in liquid N_2_, 24 hours after irradiation or 24 hours after optogenetic induction of UAS. The samples were next homogenized with pestels in RIPA buffer containing Halt™ Protease & Phosphatase Inhibitor Cocktail and left for 1 hour on ice to maximize tissue lysis. 4xLaemmli Sample Buffer + 10% 2-Mercaptoethanol was added, and samples were boiled for 10min at 95°C. Lysates were resolved by electrophoresis using Mini-PROTEAN TGX Precast Gels in Tris/Glycine/SDS Buffer. Proteins were then transferred onto 0.2μm nitrocellulose membranes using the Trans-Blot Turbo Transfer Pack, blocked in Tris-buffered saline + 0.1% Tween-20 + 5% BSA, and probed with a monoclonal anti-Xrp1 antibody^15^ (kind gift of T. Igaki). For normalization, blots were probed with Actin (Sigma-Aldrich A2103). Horseradish peroxidase (HRP)–conjugated secondary IgG antibodies were used together with the Clarity Max™ Western ECL Substrate to detect the protein bands.

### RT-qPCR

Larvae were reared at 29°C and in the dark for 72 hours after egg deposition. By incubating larvae in the light and at 29°C, RNAi expression was induced for 24 hours using the optogenetic system *Shine-Gal4*. At 96 hours after egg deposition, 50 wing discs were dissected in ice-cold PBS and quickly frozen in liquid nitrogen. Total RNA was extracted using a QIAGEN RNeasy Micro Kit according to the manufacturer’s protocol. RNA samples (2–3 μg per reaction) were treated with DNase when necessary and reverse-transcribed using SuperScript II reverse transcriptase (ThermoFisher, #18064022). The generated cDNAs were used for real-time PCR (StepOne Plus, Applied Biosystems) using Power SYBR Green PCR master mix (ThermoFisher, #A25741). Samples were normalized to *actin* and fold changes were calculated using the ΔΔCt method. Three separate biological samples were collected for each experiment, and triplicate measurements were performed. We used the following list of primers to detect specific *Xrp1* transcripts:

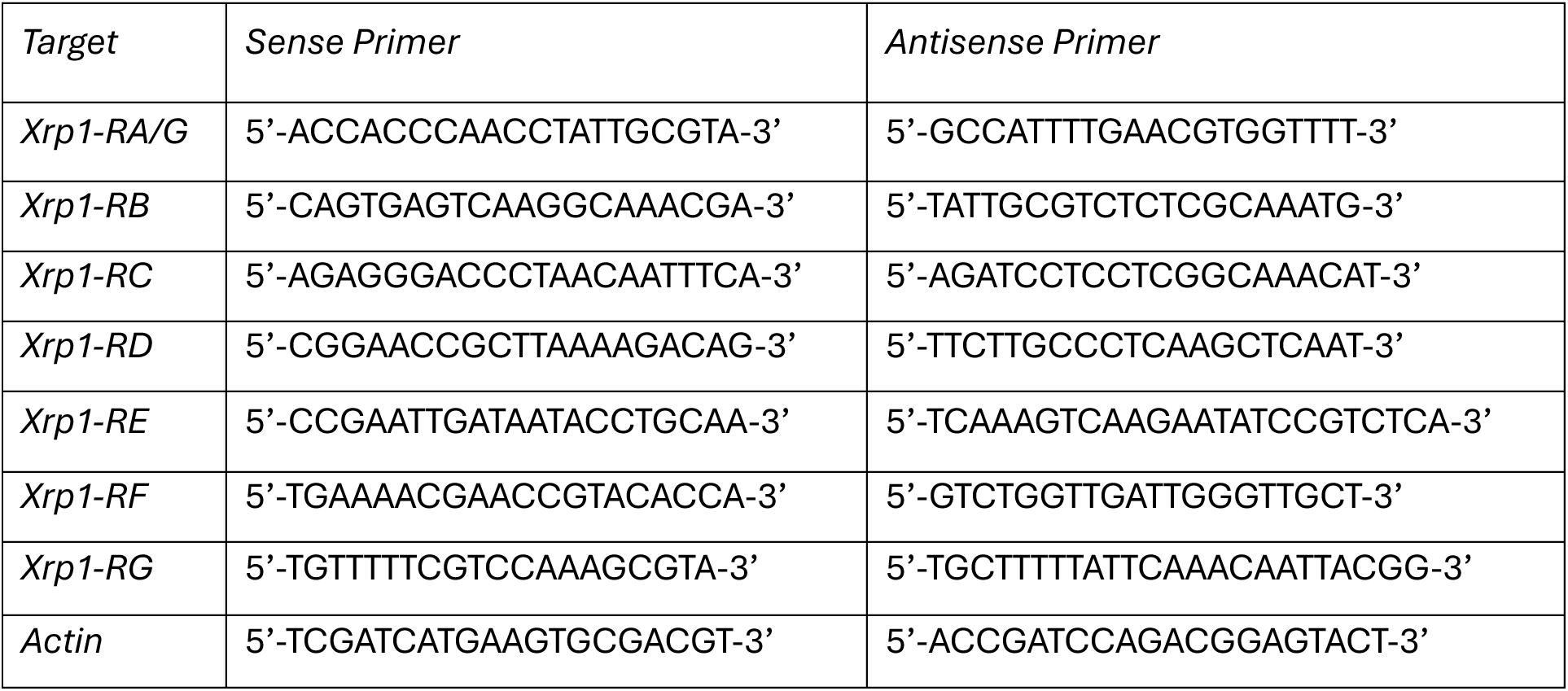

Notably, the primer set for *Xrp1-RC* can theoretically also amplify a segment in *Xrp1-RA*, *RG*, and *RB*. However, when the segment amplified from RC is around 130 bp, the amplicon derived from RA, RG and RB is expected to be around 1000 bp, which is not efficiently amplified and makes this primer set specific for RC. This was verified on the derivative melting curve plot, where a single peak was detected (Supp. Fig. 7).

---

**Supplementary Figure 1.**
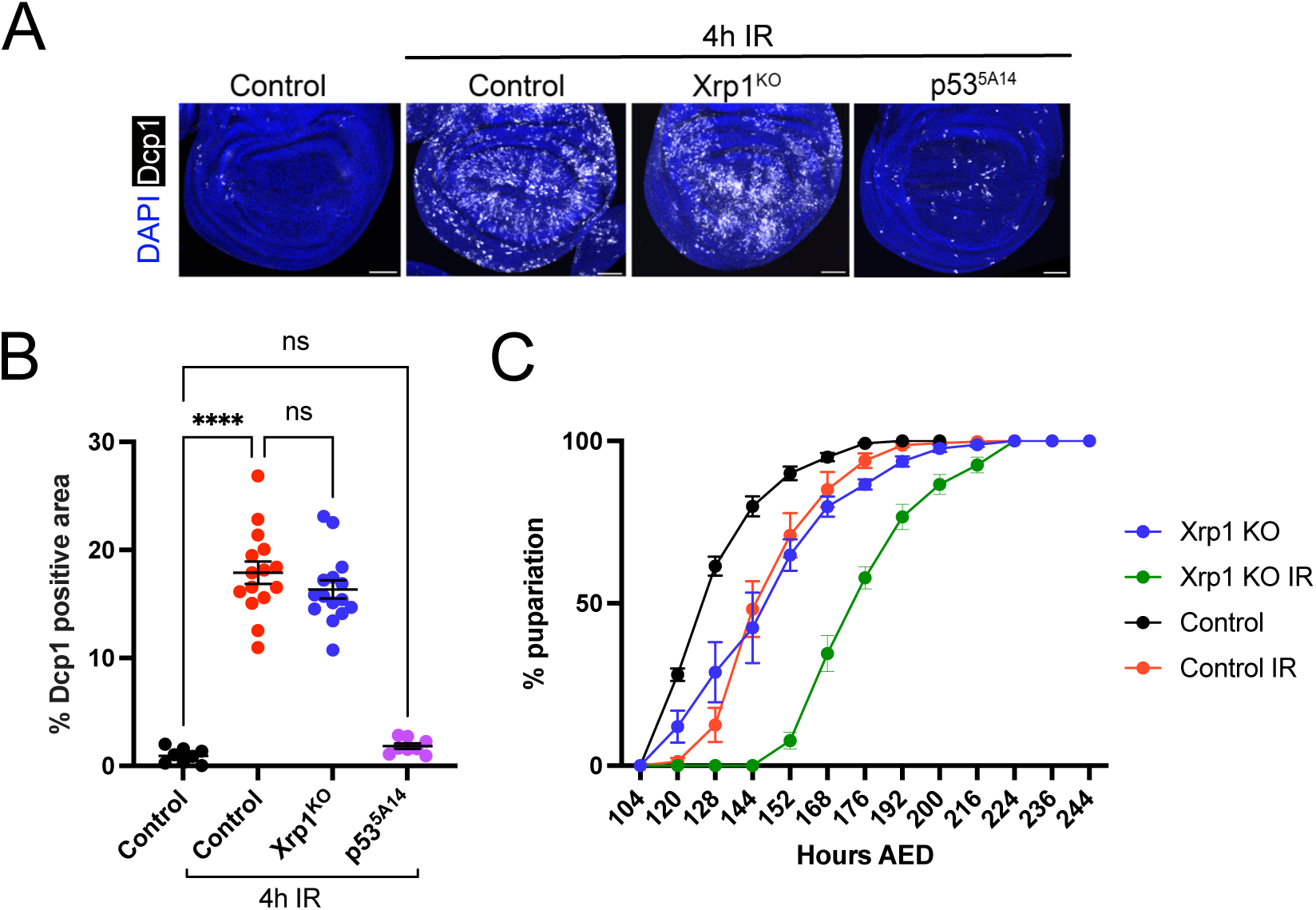
Xrp1 is not responsible for triggering apoptosis or developmental delay upon irradiation. A- Dcp1 levels used as a readout for apoptosis 4 hours after IR in an Xrp1 KO context, or p53 loss-of-function context. Wing imaginal discs were stained for DAPI and cDcp1 in control and after IR. Scale bar is 50μm. B- Quantification of Dcp1 staining in the wing imaginal discs from the conditions presented in A. Error bars indicate SEM. ****P value < 0.0001 by one-way ANOVA. ns: not significant. C- Systemic effect on developmental delay after irradiation of early L3 larvae (approximately 96hrs AED) measured by pupariation curves displaying the percentage of animals having pupariated at the indicated time points for the indicated genotypes. Error bars are SEM. AED: After egg deposition.

**Supplementary Figure 2.**
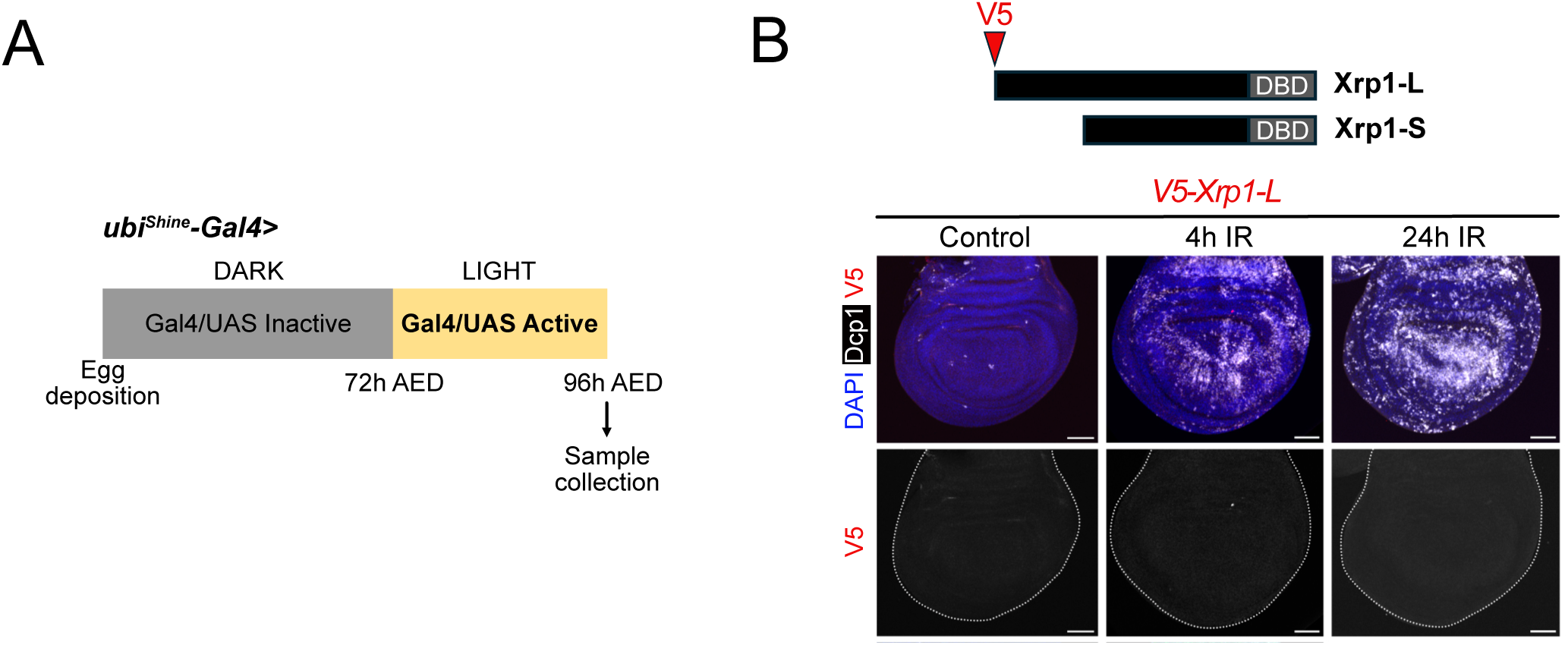
Xrp1-L is not induced upon irradiation. A- The optogenetic tool *ubi-Shine-Gal4* was used to induce RNAi expression in a controlled and non-lethal manner (see genotypes in Fig.2 B,C). Larvae were raised at 29°C, in the dark, until approximately 72 hours AED (after egg deposition). UAS expression was then induced by exposing the larvae to ambient light for 24 hours, after which their wing discs were collected. B- Representative wing imaginal discs showing the levels of a V5::Xrp1 reporter, tagging Xrp1-L specifically, 4 and 24 hours after irradiation. Max projections of discs stained for DAPI, V5, and cDcp1. Scale bar is 50μm.

**Supplementary Figure 3.**
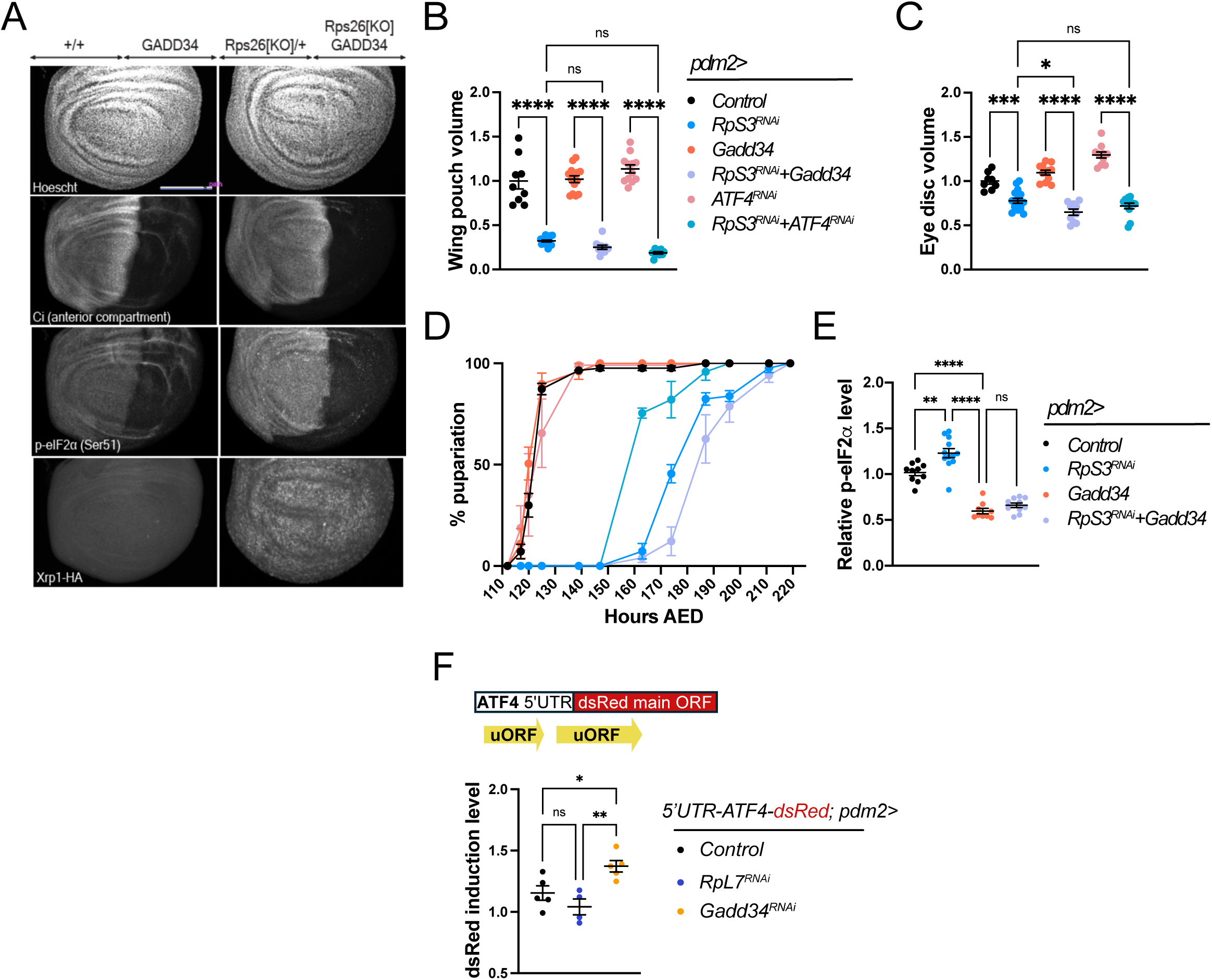
The ISR is not required for Xrp1 induction and functions downstream of a ribosomal protein deficiency. A- Representative wing discs expressing UAS-Gadd34 under the control of hh-gal4, in a control disc, or a disc lacking a copy of the *RpS26* gene (*RpS26*^KO^). Discs were stained with Hoechst, Ci (Cubitus interruptus), p-eIF2⍺, and HA, marking an Xrp1::HA reporter that tags both Xrp1 isoforms. Scale bar is 100μm. B, C, D: Autonomous and systemic effects of ISR inhibition in the context of a ribosomal protein deficiency (*RpS3-RNAi*) in the wing pouch. The genotypes are common to the three panels: B- Tissue autonomous effect measured by wing pouch volume relative to the mean value of the control genotype *pdm2>+*. ****p < 0.0001, and ns = not significant, ANOVA, error bars are SEM. C- Non-autonomous effect on remote tissue growth measured by eye disc volume relative to the mean value of the control genotype *pdm2>+*. ****p < 0.0001, *p < 0.1, and ns = not significant, ANOVA. Error bars are SEM. D- Systemic effect on developmental delay at the L/P transition, measured by pupariation curves displaying the percentage of animals having pupariated at the indicated time points. Error bars are SEM. AED: After egg deposition. E- Quantification of p-eIF2⍺ levels in the pouch, relative to the signal in the notum, showing the efficiency of Gadd34 overexpression on p-eIF2⍺ levels. ****p < 0.0001, **p<0,01, and ns = not significant, ANOVA, Error bars are SEM. F- Measurement of ATF4 translation levels using a reporter built from ATF4 5’UTR, with its uORFs, and a dsRed sequence replacing ATF4 coding region (Kang et al, 2015), upon Rp deficiency (*RpL7-RNAi*) or ISR activation (*Gadd34-RNAi*). Quantification of dsRed signal induction in the pouch, relative to the signal in the notum. **p<0,01, *p<0,1, and ns = not significant, ANOVA, Error bars are SEM.

**Supplementary Figure 4.**
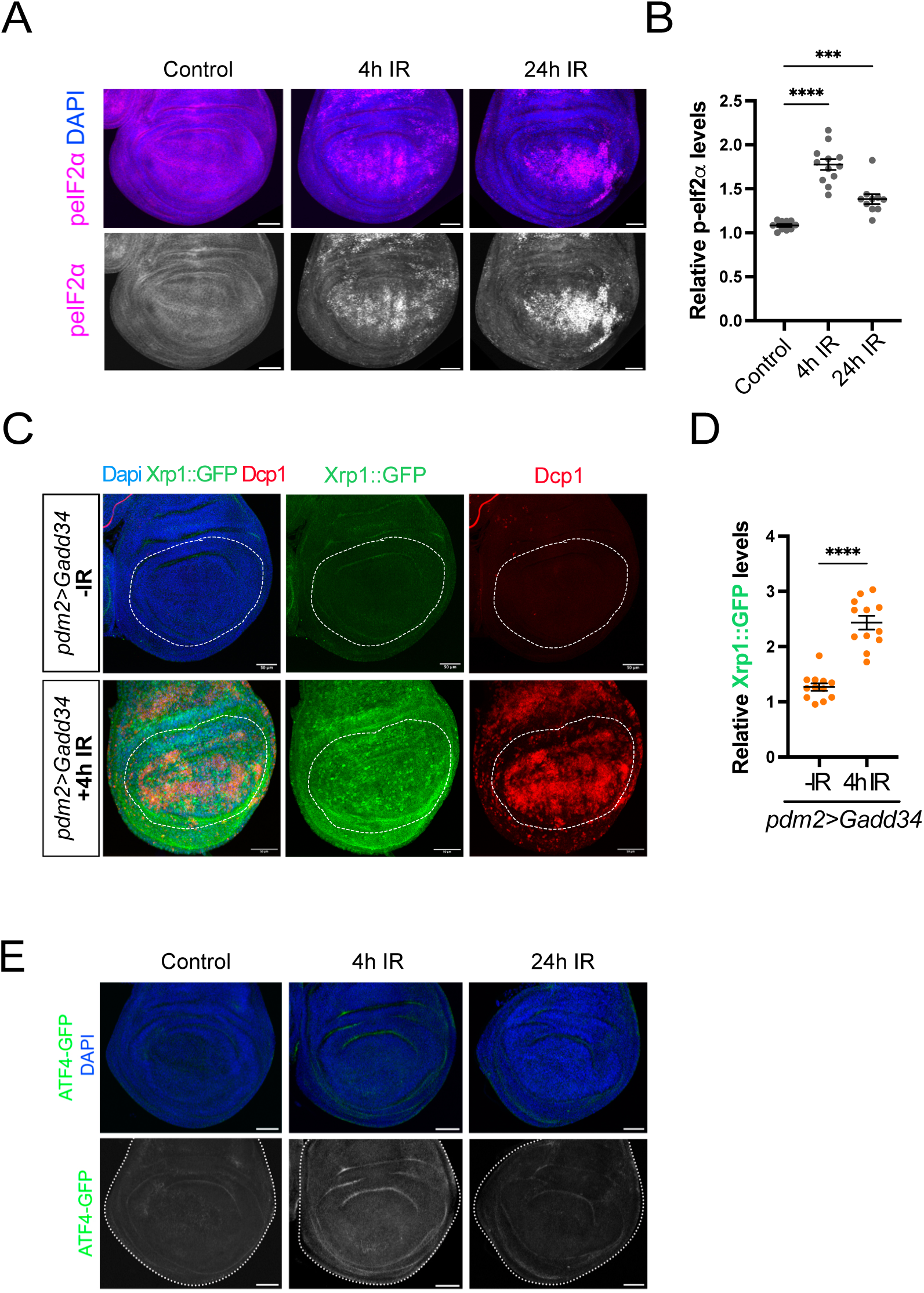
The ISR is not required for Xrp1 induction upon irradiation. A- p-eIF2⍺ accumulation 4 and 24 hours after IR. Wing discs stained for DAPI and p-eIF2⍺. Scale bar is 50μm. B- Quantification of p-eIF2⍺ levels for each condition presented in A . ****p < 0.0001 and ***p < 0.001, ANOVA, Error bars are SEM. C- Xrp1::GFP and apoptosis induction upon overexpression of GADD34, 4 hours after irradiation. Representative wing discs stained for DAPI, cDcp1, and GFP. Scale bar is 50μm. D- Quantification of Xrp1::GFP levels in the pouch, relative to the signal in the notum for the conditions presented in C. ****p < 0.0001, t-test. Error bars are SEM. E- ATF4 induction levels, as visualized with the ATF4::GFP reporter, 4 and 24 hours after IR. Wing discs stained for DAPI and GFP. Scale bar is 50μm.

**Supplementary Figure 5.**
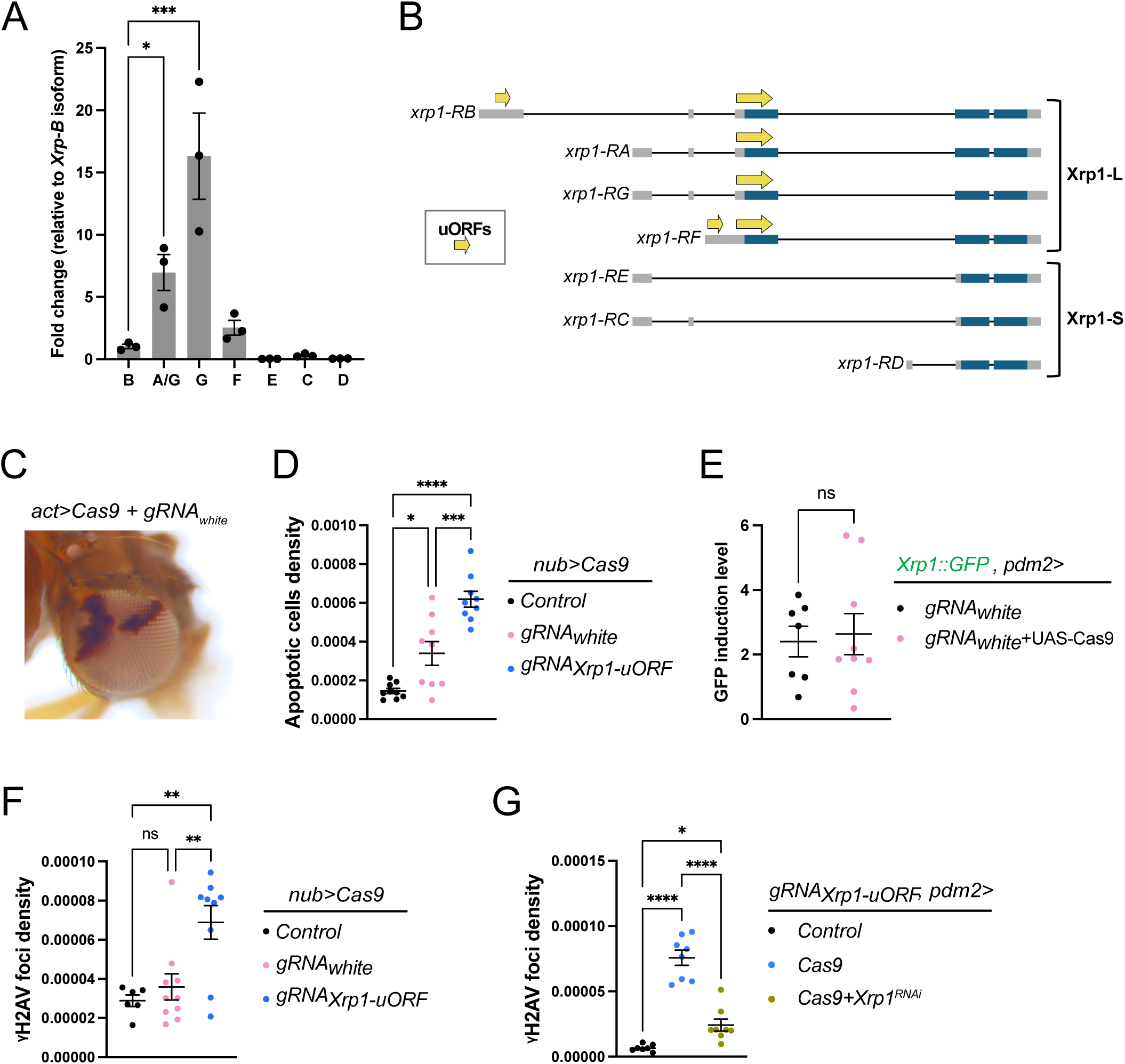
The derepression of Xrp1 is a consequence of its uORF deletion. A- Levels of *Xrp1* transcripts relative to *Xrp1-B*, measured by RT-qPCR, showing that transcripts coding for Xrp1-L are generally more abundant than those coding for Xrp1-S in physiological conditions. Values are expressed as fold changes relative to isoform B in a control genotype. ***p<0,001, and *p<0,1, ANOVA. Error bars represent SEM. The raw data used are the same as in Fig.2-C. B- Scheme representing all Xrp1-encoded transcripts. Boxes represent exons (grey are UTRs and blue are coding sequences) and fine lines are introns. The four top mRNAs give rise to Xrp1-L, while the three bottom ones translate into Xrp1-S. Only the ones encoding the long isoforms have uORFs, and in different configurations (yellow arrows). Notably, one uORF is shared by all “L- coding” isoforms and overlaps with the Xrp1-L translation start site. C- Eye of an *act>Cas9* fly, expressing guide RNAs against the *white* gene (gRNA_white_). The presence of white ommatidiae validates the guides’ efficacy. D- Quantification of apoptotic cell density in wing pouches co-expressing a Cas9 (*nub>Cas9*) and gRNAs against *white* or the Xrp1- uORF, using a cDcp1 staining. ****p<0,0001, ***p<0,001, and *p<0,1, ANOVA, Error bars are SEM. E- Quantification of Xrp1::GFP induction level in wing pouches co-expressing a Cas9 (nub>Cas9) and gRNAs against white. The GFP signal is measured as a pouch to notum ratio (t-tests, ns=non-significant, error bars are SEM). F- Quantification of the γH2Av signal, which marks DNA damage, in the wing pouches co-expressing a Cas9 (nub>Cas9) and gRNAs against *white* or the Xrp1-uORF. **p<0,01, and ns = not significant, ANOVA, Error bars are SEM. G- Quantification of the γH2Av signal, which marks DNA damage, in the wing pouch of the indicated genotypes. ****p<0,0001, and *p<0,1, ANOVA, Error bars are SEM.

**Supplementary Figure 6.**
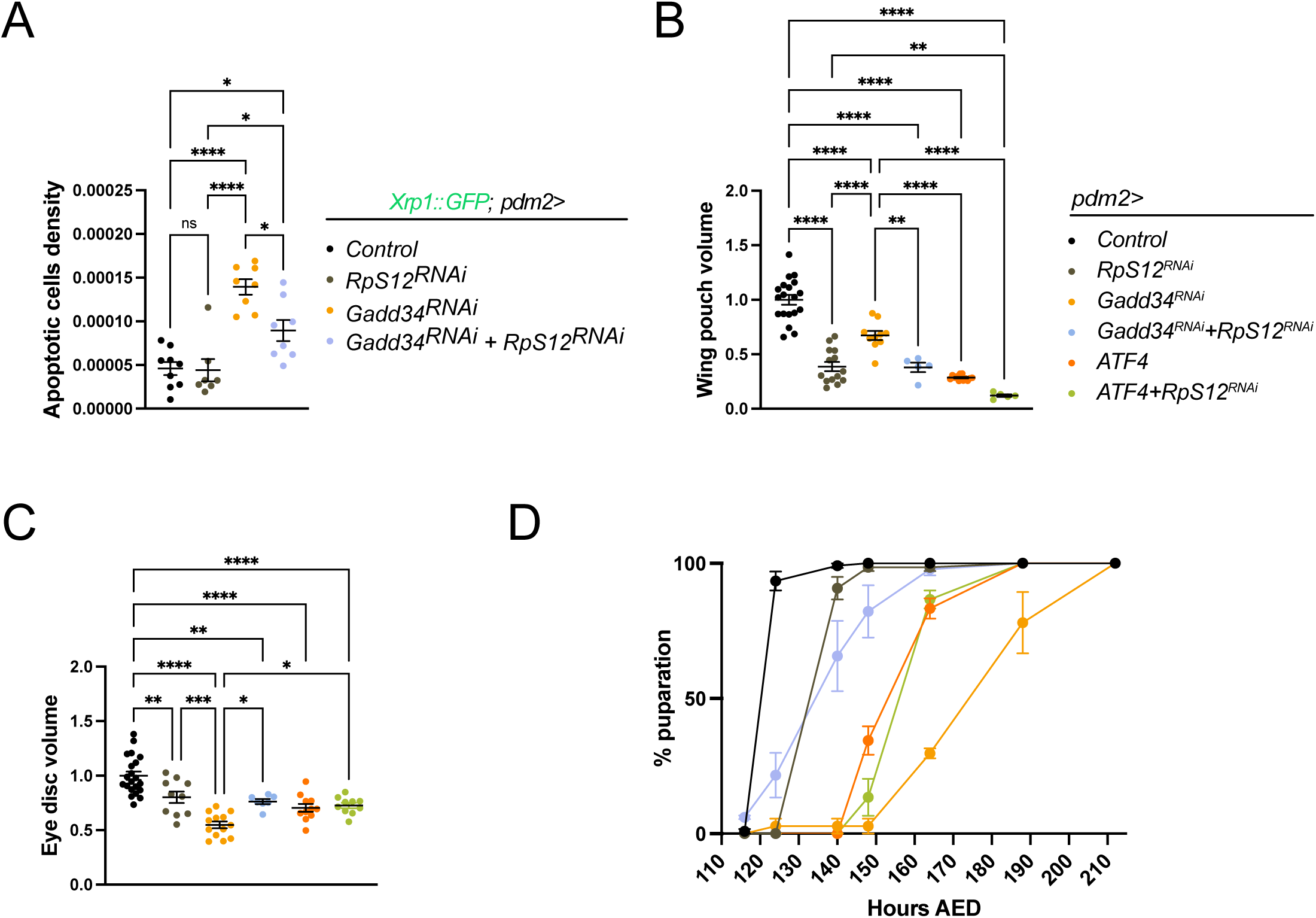
RpS12 has a role downstream of the ISR. A- Quantification of apoptotic cell density in wing pouches upon ISR activation (Gadd34-RNAi) and RpS12 inhibition (RpS12-RNAi). ****p < 0.0001, *p < 0.1, and ns = not significant, ANOVA, error bars are SEM. B, C, D: Autonomous and systemic effects of ISR activation triggered by UAS-ATF4 overexpression (bypassing uORF-mediated regulation) or UAS-Gadd34-RNAi, in the context of RpS12 knockdown in the wing pouch. The genotypes are common to the three panels: B- Tissue autonomous effect measured by wing pouch volume relative to the mean value of the control genotype *pdm2>+*. ****p < 0.0001, and **p < 0.01, ANOVA. Only significant comparisons are shown. Error bars are SEM. C- Non-autonomous effect on remote tissue growth measured by eye disc volume relative to the mean value of the control genotype *pdm2>+*. ****p < 0.0001, ***p < 0.001, **p < 0.01, and *p < 0.1, ANOVA. Only significant comparisons are shown. Error bars are SEM. D- Systemic effect on developmental delay at the L/P transition measured by pupariation curves displaying the percentage of animals having pupariated at the indicated time points. Error bars are SEM. AED: After egg deposition.

**Supplementary Figure 7.**
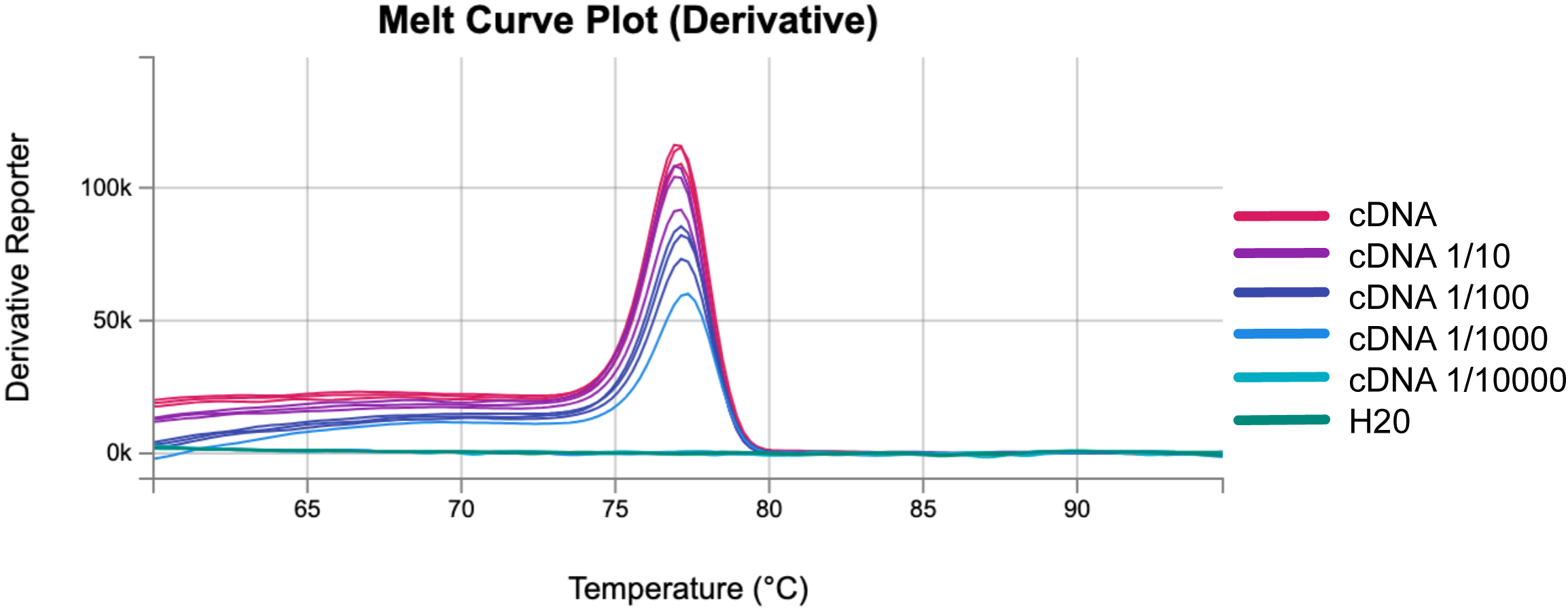
Derivative melting curve plot obtained for the primer set designed to recognize the *xrp1-RC* transcript isoform. The efficiency of the primer was tested in serial dilutions of cDNA, with concentrations ranging from 1 to 1/10 000 of the original solution.

## REFERENCES

1. Hetz, C. (2012). The unfolded protein response: controlling cell fate decisions under ER stress and beyond. Nat Rev Mol Cell Biol 13, 89–102. 10.1038/nrm3270.

2. Mitra, S., and Ryoo, H.D. (2019). The unfolded protein response in metazoan development. Journal of Cell Science 132, jcs217216. 10.1242/jcs.217216.

3. Baonza, A., Tur-Gracia, S., Pérez-Aguilera, M., and Estella, C. (2022). Regulation and coordination of the different DNA damage responses in Drosophila. Front. Cell Dev. Biol. 10, 993257. 10.3389/fcell.2022.993257.

4. Akdemir, F., Christich, A., Sogame, N., Chapo, J., and Abrams, J.M. (2007). p53 directs focused genomic responses in Drosophila. Oncogene 26, 5184–5193. 10.1038/sj.onc.1210328.

5. Costa-Mattioli, M., and Walter, P. (2020). The integrated stress response: From mechanism to disease. Science 368, eaat5314. 10.1126/science.aat5314.

6. Ryoo, H.D. (2024). The integrated stress response in metabolic adaptation. Journal of Biological Chemistry 300, 107151. 10.1016/j.jbc.2024.107151.

7. Nandakumar, S., Grmai, L., and Vasudevan, D. (2025). Emerging roles for integrated stress response signaling in homeostasis. The FEBS Journal 292, 4418–4445. 10.1111/febs.70166.

8. Baillon, L., Germani, F., Rockel, C., Hilchenbach, J., and Basler, K. (2018). Xrp1 is a transcription factor required for cell competition-driven elimination of loser cells. Scientific Reports 8, 17712–17712. 10.1038/s41598-018-36277-4.

9. Blanco, J., Cooper, J.C., and Baker, N.E. (2020). Roles of C/EBP class bZip proteins in the growth and cell competition of Rp (‘Minute’) mutants in Drosophila. eLife 9, e50535. 10.7554/eLife.50535.

10. Langton, P.F., Baumgartner, M.E., Logeay, R., and Piddini, E. (2021). Xrp1 and Irbp18 trigger a feed-forward loop of proteotoxic stress to induce the loser status. PLoS Genet 17, e1009946. 10.1371/journal.pgen.1009946.

11. Boulan, L., Andersen, D., Colombani, J., Boone, E., and Léopold, P. (2019). Inter-Organ Growth Coordination Is Mediated by the Xrp1-Dilp8 Axis in Drosophila. Developmental Cell 49, 811–818.e4. 10.1016/j.devcel.2019.03.016.

12. Baillon, L., Germani, F., Rockel, C., Hilchenbach, J., and Basler, K. (2018). Xrp1 is a transcription factor required for cell competition-driven elimination of loser cells. Scientific Reports 8, 17712–17712. 10.1038/s41598-018-36277-4.

13. Kale, A., Ji, Z., Kiparaki, M., Blanco, J., Rimesso, G., Flibotte, S., and Baker, N.E. (2018). Ribosomal Protein S12e Has a Distinct Function in Cell Competition. Developmental Cell 44, 42–55.e4. 10.1016/j.devcel.2017.12.007.

14. Tsakiri, E., Potiri, M., Kontogiannidi, K., Douka, K., Kanakousaki, K., Vavouraki, N., Loizou, M., Moulos, P., Skoulakis, E.M.C., Samiotaki, M., et al. (2026). RpS12-mediated induction of the Xrp1short isoform links ribosomal protein mutations to cell competition. Cell Reports 45, 117380. 10.1016/j.celrep.2026.117380.

15. Kakemura, B., Kanda, H., Matsumoto, R., Ueda, S., Yasuhara, S., Nagata, R., Taniguchi, K., Kondo, S., Miyoshi, K., Kobayashi, T., et al. (2026). Cellular fitness is determined by ribosomal protein S12-mediated release of a truncated Xrp1. Cell Rep 45, 117155. 10.1016/j.celrep.2026.117155.

16. Kiparaki, M., Khan, C., Folgado-Marco, V., Chuen, J., Moulos, P., and Baker, N.E. (2022). The transcription factor Xrp1 orchestrates both reduced translation and cell competition upon defective ribosome assembly or function. eLife 11. 10.7554/elife.71705.

17. Colombani, J., Andersen, D.S., and Léopold, P. (2012). Secreted peptide Dilp8 coordinates Drosophila tissue growth with developmental timing. Science (New York, N.Y.) 336, 582–585. 10.1126/science.1216689.

18. Garelli, A., Gontijo, A.M., Miguela, V., Caparros, E., and Dominguez, M. (2012). Imaginal discs secrete insulin-like peptide 8 to mediate plasticity of growth and maturation. Science (New York, N.Y.) 336, 579–582. 10.1126/science.1216735.

19. Boulan, L., Andersen, D., Colombani, J., Boone, E., and Léopold, P. (2019). Inter-Organ Growth Coordination Is Mediated by the Xrp1-Dilp8 Axis in Drosophila. Developmental Cell 49, 811–818.e4. 10.1016/j.devcel.2019.03.016.

20. Lee, J.H., Lee, E., Park, J., Kim, E., Kim, J., and Chung, J. (2003). In vivo p53 function is indispensable for DNA damage-induced apoptotic signaling in *Drosophila*. FEBS Letters 550, 5–10. 10.1016/S0014-5793(03)00771-3.

21. Wells, B.S., and Johnston, L.A. (2012). Maintenance of imaginal disc plasticity and regenerative potential in Drosophila by p53. Dev Biol 361, 263–276. 10.1016/j.ydbio.2011.10.012.

22. di Pietro, F., Herszterg, S., Huang, A., Bosveld, F., Alexandre, C., Sancéré, L., Pelletier, S., Joudat, A., Kapoor, V., Vincent, J.-P., et al. (2021). Rapid and robust optogenetic control of gene expression in Drosophila. Dev Cell 56, 3393–3404.e7. 10.1016/j.devcel.2021.11.016.

23. Recasens-Alvarez, C., Alexandre, C., Kirkpatrick, J., Nojima, H., Huels, D.J., Snijders, A.P., and Vincent, J.-P. (2021). Ribosomopathy-associated mutations cause proteotoxic stress that is alleviated by TOR inhibition. Nat Cell Biol 23, 127–135. 10.1038/s41556-020-00626-1.

24. Ochi, N., Nakamura, M., Nagata, R., Wakasa, N., Nakano, R., and Igaki, T. (2021). Cell competition is driven by Xrp1-mediated phosphorylation of eukaryotic initiation factor 2α. PLoS Genet 17, e1009958. 10.1371/journal.pgen.1009958.

25. Nagarkar-Jaiswal, S., Lee, P.-T., Campbell, M.E., Chen, K., Anguiano-Zarate, S., Cantu Gutierrez, M., Busby, T., Lin, W.-W., He, Y., Schulze, K.L., et al. (2015). A library of MiMICs allows tagging of genes and reversible, spatial and temporal knockdown of proteins in Drosophila. eLife 4, e05338. 10.7554/eLife.05338.

26. Vasudevan, D., Katow, H., Huang, H.-W., Tang, G., and Ryoo, H.D. (2022). A protein-trap allele reveals roles for *Drosophila* ATF4 in photoreceptor degeneration, oogenesis and wing development. Disease Models & Mechanisms 15, dmm049119. 10.1242/dmm.049119.

27. Kang, K., Ryoo, H.D., Park, J.-E., Yoon, J.-H., and Kang, M.-J. (2015). A Drosophila Reporter for the Translational Activation of ATF4 Marks Stressed Cells during Development. PLoS ONE 10, e0126795. 10.1371/journal.pone.0126795.

28. Kurtz, P., Jones, A.E., Tiwari, B., Link, N., Wylie, A., Tracy, C., Krämer, H., and Abrams, J.M. (2019). Drosophila p53 directs nonapoptotic programs in postmitotic tissue. Mol Biol Cell 30, 1339–1351. 10.1091/mbc.E18-12-0791.

29. Costa-Mattioli, M., and Walter, P. (2020). The integrated stress response: From mechanism to disease. Science 368, eaat5314. 10.1126/science.aat5314.

30. Dever, T.E., Ivanov, I.P., and Hinnebusch, A.G. (2023). Translational regulation by uORFs and start codon selection stringency. Genes Dev. 37, 474–489. 10.1101/gad.350752.123.

31. Brown, B., Mitra, S., Roach, F.D., Vasudevan, D., and Ryoo, H.D. (2021). The transcription factor Xrp1 is required for PERK-mediated antioxidant gene induction in Drosophila. eLife 10. 10.7554/elife.74047.

32. Xue, M., Cong, F., Zheng, W., Xu, R., Liu, X., Bao, H., Sung, Y.Y., Xi, Y., He, F., Ma, J., et al. (2023). Loss of Paip1 causes translation reduction and induces apoptotic cell death through ISR activation and Xrp1. Cell Death Discov. 9, 288. 10.1038/s41420-023-01587-8.

33. Ji, Z., Kiparaki, M., Folgado, V., Kumar, A., Blanco, J., Rimesso, G., Chuen, J., Liu, Y., Zheng, D., and Baker, N.E. (2019). Drosophila RpS12 controls translation, growth, and cell competition through Xrp1. PLoS Genet 15, e1008513. 10.1371/journal.pgen.1008513.

34. Lee, C.-H., Kiparaki, M., Blanco, J., Folgado, V., Ji, Z., Kumar, A., Rimesso, G., and Baker, N.E. (2018). A Regulatory Response to Ribosomal Protein Mutations Controls Translation, Growth, and Cell Competition. Developmental Cell 46, 456–469.e4. 10.1016/j.devcel.2018.07.003.

35. Anger, A.M., Armache, J.-P., Berninghausen, O., Habeck, M., Subklewe, M., Wilson, D.N., and Beckmann, R. (2013). Structures of the human and Drosophila 80S ribosome. Nature 497, 80–85. 10.1038/nature12104.

36. Martín-Villanueva, S., Galmozzi, C.V., Ruger-Herreros, C., Kressler, D., and De La Cruz, J. (2024). The Beak of Eukaryotic Ribosomes: Life, Work and Miracles. Biomolecules 14, 882. 10.3390/biom14070882.

37. Brito Querido, J., Sokabe, M., Díaz-López, I., Gordiyenko, Y., Fraser, C.S., and Ramakrishnan, V. (2024). The structure of a human translation initiation complex reveals two independent roles for the helicase eIF4A. Nat Struct Mol Biol 31, 455–464. 10.1038/s41594-023-01196-0.

38. Walsh, K., Katow, H., Junn, H., Vasudevan, D., Dieterich, C., and Ryoo, H.D. (2025). 4EHP and NELF-E regulate physiological ATF4 induction and proteostasis in disease models of Drosophila. Nat Commun 17, 626. 10.1038/s41467-025-67357-5.

39. Zhang, B., Mehrotra, S., Ng, W.L., and Calvi, B.R. (2014). Low levels of p53 protein and chromatin silencing of p53 target genes repress apoptosis in Drosophila endocycling cells. PLoS Genet 10, e1004581. 10.1371/journal.pgen.1004581.

40. Di Pietro, F., Herszterg, S., Huang, A., Bosveld, F., Alexandre, C., Sancéré, L., Pelletier, S., Joudat, A., Kapoor, V., Vincent, J.-P., et al. (2021). Rapid and robust optogenetic control of gene expression in Drosophila. Developmental Cell 56, 3393–3404.e7. 10.1016/j.devcel.2021.11.016.

41. Poernbacher, I., Crossman, S., Kurth, J., Nojima, H., Baena-Lopez, A., Alexandre, C., and Vincent, J.-P. (2019). Lessons in genome engineering: opportunities, tools and pitfalls. Preprint at Genetics, 10.1101/710871 10.1101/710871.

42. Baena-Lopez, L.A., Alexandre, C., Mitchell, A., Pasakarnis, L., and Vincent, J.-P. (2013). Accelerated homologous recombination and subsequent genome modification in Drosophila. Development 140, 4818–4825. 10.1242/dev.100933.

